# A toolkit for transposon libraries and functional genomics in intestinal Bacteroidales

**DOI:** 10.1101/2025.10.10.681549

**Authors:** Carlos Geert Pieter Voogdt, Indra Roux, Katharina Müller, Nicolai Karcher, Afonso Martins Bravo, Lajos Kalmar, Vallo Varik, Jacob Bobonis, Georg Zeller, Michael Zimmermann, Kiran Raosaheb Patil, Athanasios Typas

## Abstract

Members of the order Bacteroidales include some of the most prevalent and abundant bacterial species in the healthy human gut microbiota. Yet, most of the functions encoded in their genomes remain poorly characterized, limiting our understanding of the different roles they play in the human gut microbiome. Towards addressing this gap, we developed tools and methods for genome-wide transposon mutagenesis in Bacteroidales, including broad-range transposon vectors with several antibiotic selection markers, a dual conjugation-cloning donor strain, and protocols for convenient library generation in liquid media. We then created saturated, barcoded, insertion mutant libraries in the type strains of three key representatives of the main genera within Bacteroidales: *Bacteroides uniformis* (ATCC 8492), *Phocaeicola vulgatus* (ATCC 8482) and *Parabacteroides merdae* (ATCC 43184). Based on the dense transposon insertion profiles and a workflow for comparing essentialomes across species, we identified 275 core essential genes shared across the three species, and 163 species-specific essential genes, some of which could be explained by functional redundancy and alternative metabolic pathways. We further identified essential non-protein coding elements and essential protein domains with known and unknown functions. Finally, using insertion directionality bias, we could map potential toxic modalities in the three genomes, including toxin-antitoxin pairs, mobile elements encoding toxic products and enzymes leading to toxic metabolic intermediates. Overall, the tools, workflows and genome-wide resources reported here expand the experimental repertoire for characterizing genes in key bacteria of the human gut microbiome, and pave the way for the establishment of similar genetic toolkits for other gut bacteria.

## Introduction

Advances in sequencing technologies have revealed an unprecedented taxonomic and genomic diversity within microbial ecosystems^1–6^. Yet, this enormous diversity of microbial gene sequences is also highlighting how little we know about their encoded functions. Even in well-studied model organisms, such as *Escherichia coli* and *Bacillus subtilis,* up to a third of their genes remain poorly characterized^7,8^. The knowledge gap is much larger for the many non-model microbes found in diverse ecosystems. High-throughput reverse genetics, which systematically link genes to each other and to phenotypes, offers a powerful strategy to close this gap in microbial gene function annotation^9–14^. However, applying such approaches to non-model microbes requires efficient genome-wide genetic tools.

Random transposon mutagenesis has been used for decades to generate mutant libraries in bacteria. By inserting at random genomic locations, transposons disrupt genes and create loss-of-function mutant libraries without the need for targeted genetic tools. While originally used for forward genetic screens, the declining cost of sequencing and the incorporation of random DNA barcodes into the transposon^15,16^, have transformed transposon mutant libraries into powerful resources for systematic genotype-to-phenotype mapping^17–19^. Libraries containing insertion mutants for thousands of genes can be assessed in hundreds of conditions, such as nutrients, xenobiotics or host colonization, to reveal genes important for fitness in specific environments and to infer functional links between genes with similar phenotypic profiles^14,20–22^. Furthermore, high density transposon mutagenesis also enables the assessment of gene essentiality. In dense libraries, genes and other genetic features intolerant to disruptive insertions can be identified as essential, revealing core functions critical for growth and survival^23–25^. Gene essentiality can be context-dependent, changing with the environment and/or genetic background^9,26,27^. Thus, identifying and comparing essential genes across species can both highlight shared central functions and help to understand species-specific physiology.

One of the most extensively studied microbial ecosystems is the human intestinal microbiota. This ecosystem contains 200-500 bacterial species, considerably varying across individuals at the strain-level, and therefore also, at the gene level^28,29^. Since most of these species are phylogenetically distant from classical model bacteria, their genomes contain a large fraction of uncharacterized genes. To expedite their functional interrogation, transposon mutant libraries have been made in several human gut bacterial species, such as *Enterococcus faecalis*^30^, *Akkermansia muciniphila*^31^, *Clostridioides difficile*^32^, *Bifidobacterium breve*^22^, *Lactobacillus casei*^33^, and several members of the order Bacteroidales^20,21,34–38^. Part of one of the two major phyla found in the human gut microbiome (Bacteroidota), Bacteroidales contain some of the most prevalent and abundant species of the healthy human gut microbiota. These species are mainly part of three genera: *Bacteroides*, *Phocaeicola* and *Parabacteroides*^39^, some of which have been associated with health benefits such as the production of short chain fatty acids through fermentation of dietary complex carbohydrates^40–42^.

While dense transposon libraries have been generated in Bacteroidales species, these have almost exclusively focused on the *Bacteroides* genus and in particular two of its species, *B. thetaiotomicron* and *B. fragilis*. Many other prevalent and abundant Bacteroidales species of the healthy human gut microbiota remain genetically unexplored. Transposon mutagenesis is well suited to address these gaps, but generating dense libraries across diverse gut Bacteroidales species and strains requires versatile vectors and easy-to-implement methods and analysis pipelines. To meet this need, we developed a set of broad-range, barcoded transposon vectors and a multi-purpose *E. coli* conjugation donor for the generation of high-density barcoded transposon insertion libraries in multiple Bacteroidales species and strains. We used these genetic tools to generate saturated, barcoded libraries in the type strains of three prominent gut species, *B. uniformis*, *Parabacteroides merdae* and *Phocaeicola vulgatus,* representatives of the three main genera within the order. In addition to conventional library construction via outgrowth on solid media, we established a liquid-based protocol, which is considerably easier and more scalable in confined anaerobic workstations. We demonstrated the utility of these dense libraries through comparative analyses that revealed conserved and species-specific essential genes, essential protein domains, and toxic genetic modalities across the three species.

## Results

### Novel tools for transposon mutagenesis in Bacteroidales

We set out to create a broad-range barcoded transposon vector able to mutagenize multiple species of Bacteroidales with high efficiency. We used the pSAM-bt plasmid^35^ as the basis, and changed several of its features. The *B. thetaiotaomicron rpoD* promoter that drives expression of the Himar1-C9 transposase was replaced by the strong constitutive *B. fragilis* phage promoter P_BfP1E6_, which is active in different *Bacteroides* species^43^. The native *B. thetaiotaomicron* promoter driving *ermG* expression (erythromycin resistance) was replaced by the hybrid *cepA* promoter^44^, as this promoter was previously shown to limit insertional bias in *B. thetaiotaomicron* transposon mutagenesis^21^. Further, we removed the transcriptional terminator in the pSAM-bt transposon to avoid polar mutations in operons, where early termination can block transcription of downstream genes. In its place, we inserted a random sequence flanked by BsmBI restriction sites to enable plasmid barcoding through Golden Gate cloning^45,46^. The barcode entry site is flanked by primer binding sites for barcode sequencing (BarSeq^16^). As resistance to erythromycin is common among various species of Bacteroidales^47^, we also created vectors in which the transposon encodes *catP* or *tetQ*, instead of *ermG*^48^, providing resistance to chloramphenicol or tetracycline, respectively (Fig. 1A, see supplementary file 1 for plasmid features).

**Fig. 1:**
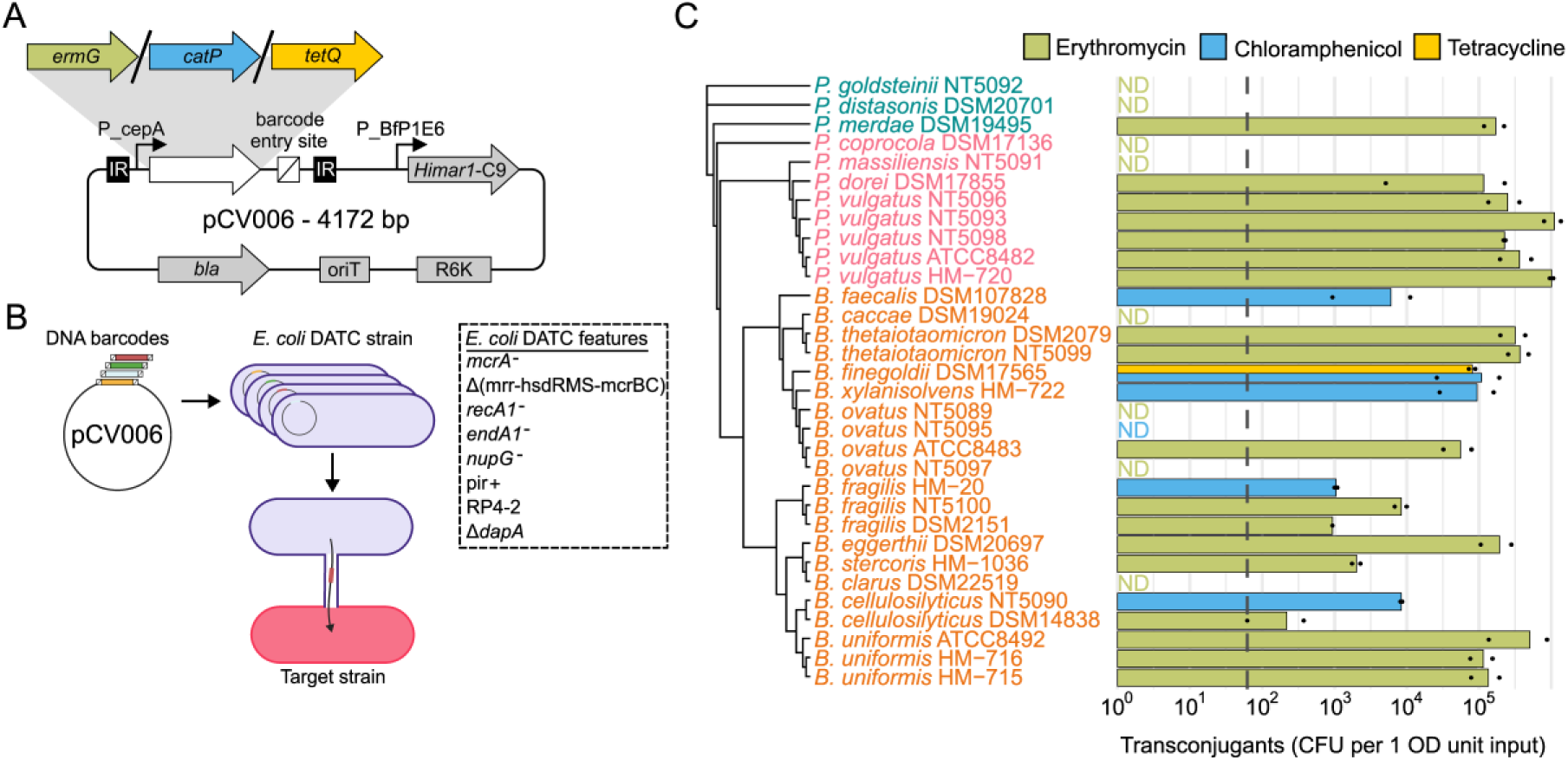
Optimized barcoded transposon mutagenesis in Bacteroidales. **A)** A broad-host-range transposon vector series for Bacteroidales, carrying different selection markers and a BsmBI based cloning site (Golden Gate) for barcoding. The barcoded plasmid pools are also available. Expression of the transposase and selection markers are placed under conserved and constitutive Bacteroidales promoters. IR: inverted repeat, *bla*: beta-lactamase, *oriT*: origin of transfer, R6K: origin of replication, *himar1*-C9: transposase (mariner), *ermG*: rRNA adenine N-6-methyltransferase (erythromycin resistance), *catP*: chloramphenicol acetyltransferase, *tetQ*: tetracycline resistance protein. **B)** A universal donor strain for efficient broad-host range conjugation. The *E. coli* strain DATC can be used both for plasmid barcoding/cloning and conjugation. The schematic highlights some of its genomic features that facilitate both processes. **C)** The new universal donor strain and broad-host range vector series allow for high conjugation efficiency across three Bacteroidales genera. Conjugation efficiency of 32 recipient strains of *Parabacteroides* (green), *Phocaeicola* (pink) and *Bacteroides* (orange), as the number (Colony Forming Units) of transconjugants per 1 OD unit of input cells (which is 1 ml of saturated culture of OD = 1). ND: no transconjugants detected. Dashed line denotes the limit of detection. Bars indicate the mean CFU count of two biological replicates (replicates shown as black dots) on mGAM agar plates containing appropriate antibiotics. Strains are grouped by phylogeny according to a neighbor-joining tree built on whole genome average nucleotide identity approximated with Mash version 2.3^109^.

High transformation efficiency is necessary for generating diverse collections of barcoded transposon plasmids. The transformation efficiency of pCV006 (*ermG* version) was very low in the widely used *E. coli* S17-1 conjugation donor, but much higher in the *E. coli* cloning strain EC100D*pir*+ (Extended Data Fig. 1A). Therefore, we engineered EC100D*pir*+ to create a dual cloning-conjugation strain^49^. For this, we used phage P1*vir* to transduce the EC100D*pir*+ cloning strain with a P1 lysate from *E. coli* MFD*pir*+^50^, in which the RP4 conjugation locus is flanked by neomycin and apramycin resistance genes, allowing for selection in both antibiotics. We also deleted *dapA* from the chromosome to make the conjugation donor auxotrophic for diaminopimelic acid (DAP), providing a counter-selectable marker. The resulting strain, DATC (DAP Auxotroph Transformation Conjugation donor^49^), has useful genomic features for both cloning and conjugation (Fig. 1B). Importantly, it exhibited high transformation efficiency with pCV006, as its EC100D*pir*+ parent (Extended Data Fig. 1A).

We tested the ability of our new transposon vectors and the DATC *E. coli* donor strain to mutagenize a panel of 32 Bacteroidales strains (21 species from 3 genera). We conjugated either the *ermG*, *catP* or *tetQ* versions of the vector into each of the 32 strains (Fig. 1C), after determining their resistance to erythromycin, chloramphenicol, and tetracycline (Extended Data Fig. 1B). We detected substantial transposon insertion mutagenesis in ∼72% (23/32) of the tested recipient strains and in representatives of all 3 genera. The few recipient strains for which we failed to detect transconjugants were spread across the phylogenetic tree, indicating that endogenous strain-specific defenses against conjugation or foreign DNA elements were likely the underlying cause, rather than incompatibility of our vector with specific species. Overall, we generated a set of transposon vectors and a dual cloning-conjugation strain that are useful for transposon mutagenesis in a broad range of Bacteroidales species and strains.

### Generation of saturated barcoded transposon libraries in three genera

We used our mutagenesis system to generate high-density, barcoded libraries in the type strains of *Bacteroides uniformis* (ATCC 8492), *Phocaeicola vulgatus* (ATCC 8482) and *Parabacteroides merdae* (ATCC 43184). We first sequenced the three strains and obtained closed, single-contig genome assemblies, which allowed us to confidently map the transposon insertions. Additionally, to facilitate comparability in downstream analyses, all genomes were annotated using the mettannotator suite^51^. To render our libraries amenable for barcoded sequencing (BarSeq^16^) and further library mutliplexing^38^, we inserted DNA barcodes of 25 random nucleotides into pCV006 using Golden Gate cloning (Methods), and a four nucleotide, library-specific index in front of the barcode. Following overnight conjugation of the DATC donor carrying the barcoded pCV006 with *B. uniformis*, *P. vulgatus* or *P. merdae* under aerobic conditions (Fig. 2A), we plated the mutant libraries on more than 25 large (145 mm) petri dishes containing Modified Gifu Anaerobic Medium (mGAM) with erythromycin and incubated these until single colonies were detectable. While outgrowth on solid media is a standard step in transposon library construction, it becomes cumbersome in confined anaerobic chambers, since it limits handling and available space. As an alternative, we conducted the outgrowth of the mutant libraries for 25-28 generations in a single bottle containing 100 ml selective mGAM liquid growth medium – this number of generations approximates the growth required for a single cell to form a visible colony. We cryo-preserved both the solid- and liquid-generated libraries, and used a single library aliquot for transposon insertion sequencing (TnSeq)^52^ using a 2-step semi random PCR procedure^53^ (Methods). To map the transposon insertion sites to the three genomes, we used TnSeeker (Methods), an in-lab developed software that maps raw TnSeq reads onto the assembled genome, and creates a list of mutants for each library with positional, directional, and barcode information (Fig. 2A).

**Fig. 2:**
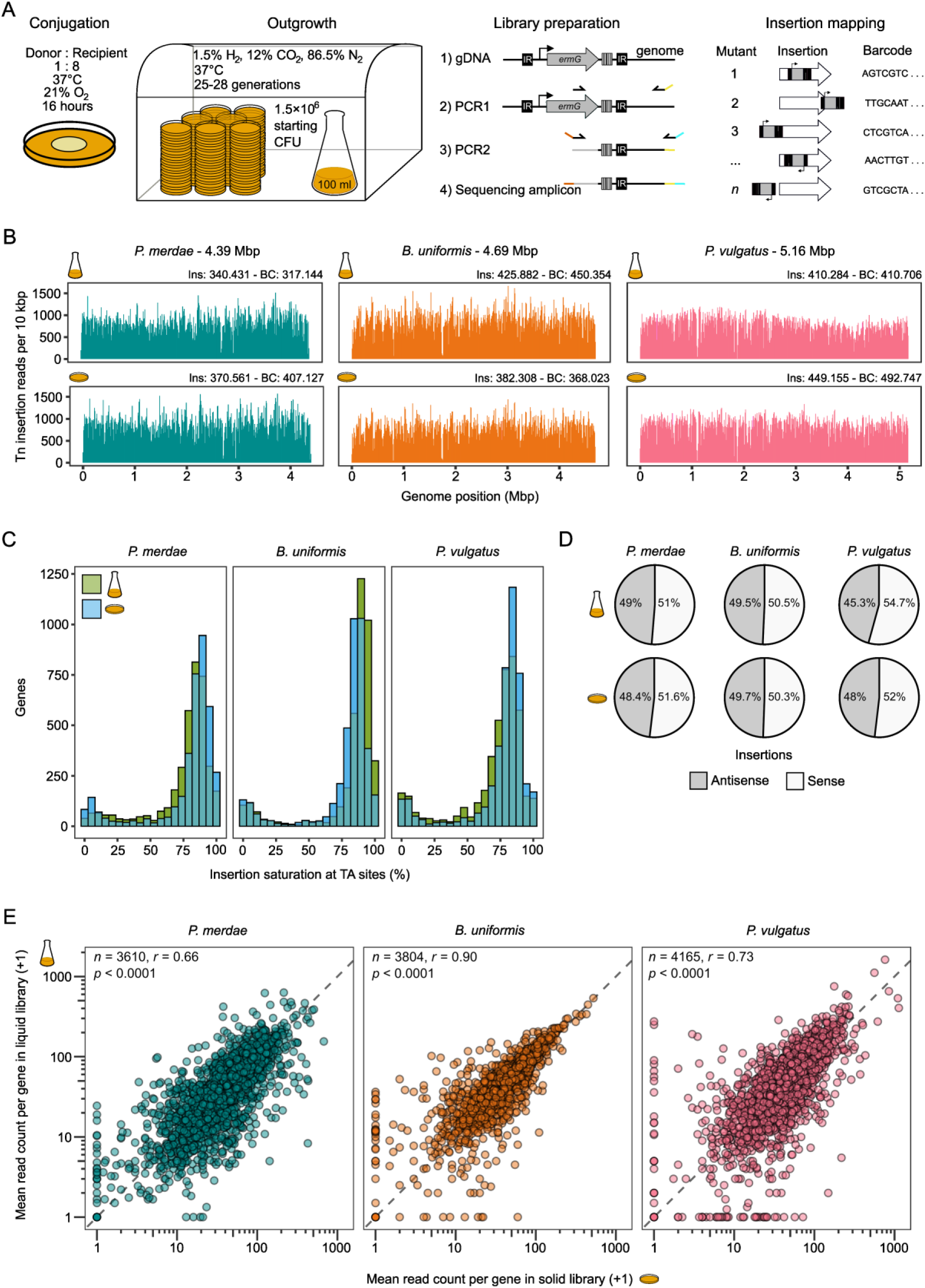
Saturated barcoded transposon libraries in three Bacteroidales genera. **A)** Schematic of a simplified protocol to create transposon insertion libraries in Bacteroidales species. Following overnight conjugation on mGAM plates in aerobic conditions, ∼1.5 million transconjugants are inoculated in liquid media (simplified method that improves handling) or plated on more than 25 large petri dishes (solid/conventional method). After 25–28 generations of outgrowth, the liquid culture or colonies scraped from all plates are used for cryostocking and sequencing library preparation, which involves DNA extraction followed by two sequential PCRs to generate amplicons ready for sequencing. Insertion and barcode mapping are done using TnSeeker (Methods). **B)** Dense and unbiased transposon insertions into the genome of *P. merdae*, *B. uniformis* and *P. vulgatus* type strains. Transposon insertion coverage across the genome is shown as the sum of insertion reads per 10 kbp. Genome size, number of unique insertion positions (Ins) and unique barcodes (BC) are indicated above each plot. **C)** Most non-essential genes are highly saturated with insertions in all three species, independent of whether the conventional solid or simplified liquid protocol was used. Genes are binned based on the insertion saturation level of TA dinucleotide sites which is the number of TA sites with insertions as a percentage of all TA sites within a gene. **D)** Solid- and liquid-based libraries show no directionality bias of insertions in the sense (coding) or antisense strand of the targeted gene. **E)** Mutant abundance is similar in solid- and liquid-generated libraries. Shown is the mean read count of all mutants per gene. One count was added to the reads to enable visualization on a log scale. *n* = number of genes analyzed, *r* = Pearson correlation coefficient, *p* = *p*-value (two-sided).

We identified over 340,000 transposon insertion sites in all six libraries (three species, solid or liquid outgrowth), spread evenly across the genome without insertion bias (Fig. 2B). Barcode analysis revealed that some sites were represented by more than one mutant, and after barcode filtering, we retained between 317,144 and 492,747 uniquely mapping barcodes per library (see Methods). No major differences were observed in the abundance of unique barcodes between species or outgrowth procedure (liquid vs. solid) and across all libraries, with the median number of barcodes per gene being between 54 and 82, and 75% of genes having at least 19 unique barcodes (Extended Data Fig. 2A). Multiple unique barcodes per gene represent independent mutants for this gene, and provide multiple biological replicates in BarSeq experiments, increasing the statistical power for assessing gene fitness^16^.

Further characterization of the libraries showed that more than 90% of insertions occurred at TA dinucleotides, as expected for Himar Mariner transposases (Extended Data Fig. 2B), and that the proportion of TA sites with insertions was similar across intragenic, intergenic and annotated non-protein-coding elements, indicating no apparent bias among different genomic elements (Extended Data Fig. 2C). Our libraries also exhibited high insertion saturation, with an average of 76% of genes across the six libraries having more than 75% of their TA sites disrupted by a transposon (Fig. 2C). In line with previous reports on *B. thetaiotaomicron*^21^, we observed no directional bias in the *B. uniformis* and *P. merdae* libraries and only a very slight bias in the *P. vulgatus* libraries towards insertions in the sense strand of the targeted gene (Fig. 2D). This suggests that the P*_cepA_*-*ermG* promoter-gene combination is well-suited to generate libraries in Bacteroidales species with little to no overall directional bias.

To assess whether the liquid outgrowth protocol affected the library composition, we compared the relative abundances of disrupted genes between liquid- and solid-generated libraries in the three species, approximating gene abundance by the mean read count of its mutants. Although differences between the libraries were observed (see also essentiality analysis later), mostly for genes with low read counts, the Pearson correlation coefficients were high in all three species (0.66-0.90; Fig. 2E). Thus, as outgrowth in liquid media did not compromise library diversity or introduce systematic bias, we suggest that this liquid protocol can serve as a robust, easy, and cost-effective alternative for generating libraries in gut microbes.

### Conservation and species-specificity of gene essentiality in Bacteroidales

We used our transposon libraries to systematically map and compare gene essentiality across *B. uniformis*, *P. vulgatus* and *P. merdae*. The saturation of our libraries made them ideal for predicting gene essentiality with high confidence. We analyzed the TnSeq data using the software package TRANSIT to identify essential genes via a Hidden Markov Model (HMM) and a Bayesian/Gumbel model^54^. To arrive at a single essentiality call per gene, we first consolidated high confidence calls from the HMM and Gumbel methods per library (Extended Data Fig. 3A) and then further consolidated these between the liquid and solid libraries (Extended Data Fig. 3B&C). All essentiality data, including the HMM and Gumbel output, of the three species is provided in supplementary table 1.

About 9% of the coding genes in each of the three species were classified as essential (Fig. 3A). This proportion is similar to previous findings in other Bacteroidales, including *B. thetaiotaomicron* (6.7%^35^ and 7.9%^21^) and *Bacteroides fragilis* (12.7%^37^). Similar ranges have been reported in model bacteria, such as *E. coli* (8.3%^25^), *Bacillus subtilis* (6.1%^10^), *Caulobacter crescentus* (12.2%^24^), and several Enterobacteriaceae species (6–9.1%^55^). Importantly, for most genes the essentiality calls agreed between solid and liquid libraries (Extended Data Fig. 3D). In *P. merdae*, *B. uniformis* and *P. vulgatus*, 11.2%, 3.1% and 7.1% of the essential genes respectively, were flagged as being essential in only the solid or liquid library (Fig. 3A). Most of these differential calls were due to low confidence by the HMM and Gumbel method in one of the libraries (Supplementary Table 1), but in some cases biological reasons may explain the differences. For example, pyrimidine biosynthesis genes in *P. merdae* were essential only on solid media (Extended Data Fig. 3E), possibly because metabolite cross-feeding is limited on agar plates, but can occur in liquid culture, rescuing the auxotrophy of these mutants.

**Fig 3.**
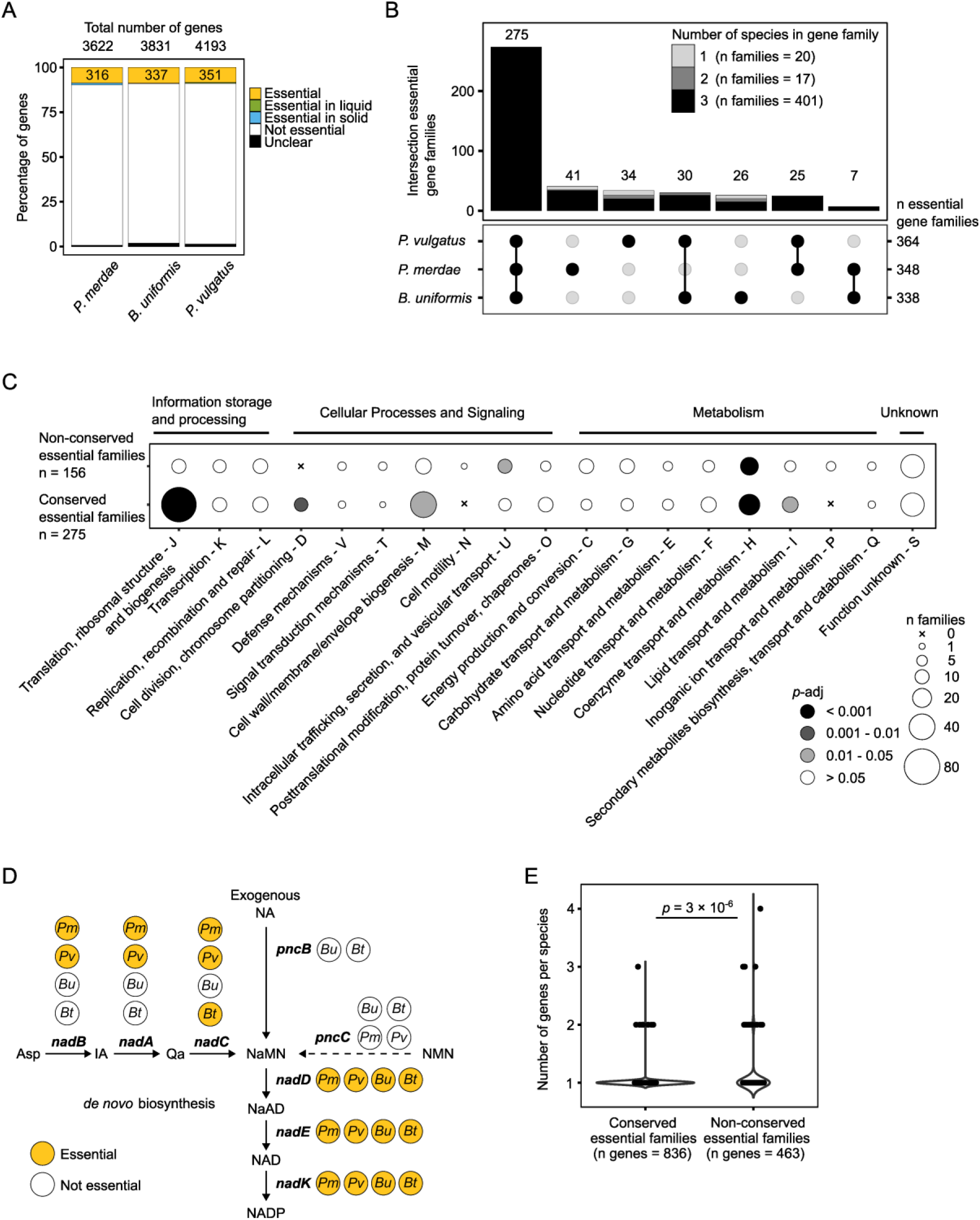
Gene essentiality across three species of Bacteroidales. **A)** Less than 10% of all genes are required for growth in rich media (mGAM) in all three species. At most, ∼1.1% (*P. merdae*) of genes are only essential in either liquid or solid growth conditions. Essentiality calling is based on transposon insertion coverage per gene (Methods). **B)** Essential genes are largely conserved across the three genera with evidence of species-specific bypassing of gene essentiality. Over 60% of essential gene families are conserved across all three species, and over 75% in at least two species. Shown are essential gene families as defined by Cluster of Orthologous Groups (COGs) at the Bacteroidia taxonomic level (class). Bar shading reflects the number of tested species containing the COG. For example, among the 41 families that are essential only in *P. merdae*, most also include non-essential orthologs in the other two species. **C)** Essential genes are enriched in core cellular processes. Functional enrichment analysis of conserved and non-conserved essential gene families from the three species using Cluster of Orthologous Groups (COG) functional categorization (indicated by capital letters). Dot shading represents the adjusted *p*-value (Benjamini–Hochberg correction) of a one-sided Fisher’s exact test (testing for enrichment). Of the 163 non-conserved essential gene families, 7 lacked COG annotation and were excluded from the analysis. COG categories without essential gene families are not shown. **D)** The *de novo* pathway for NAD biosynthesis is essential in *P. merdae* and *P. vulgatus*, but not in *B. uniformis* and *B. thetaiotaomicron* (based on prior essential analysis data^21^), which encode a salvage pathway from exogenous nicotinic acid (NA). The dashed line marks another putative salvage pathway through *pncC,* yet this does not enable bypassing of essentiality. Asp: Aspartate; IA: iminoaspartate; Qa: quinolinic acid; NaMN: nicotinate mononucleotide; NaAD: nicotinate adenine dinucleotide; NMN, nicotinamide mononucleotide; NAD(P): nicotinamide adenine dinucleotide (phosphate). **E)** Non-conserved essential genes are more frequently found in higher copy numbers than conserved essential genes across the three genera. Therefore, redundancy is likely a common mechanism for bypassing essentiality. Shown are the number of genes within conserved essential families across all three species (n = 275) against those within non-conserved essential families (n = 163). Statistical significance was assessed using a two-sided Wilcoxon rank-sum test. For specific examples see Extended Data. Fig. 4D.

We next investigated the conservation and divergence of gene essentiality among the three species. Genes were grouped into families using the Cluster of Orthologous Groups (COGs) at the Bacteroidia (class) taxonomic level from the EggNOG 5.0 database^56^, which yielded 452 families containing at least one essential gene. Among these, 63 families lacked representation from one or two of the species. Such absences could reflect genuine species-specific differences, or alternatively, cases where orthologous genes were assigned to different families. To distinguish between these possibilities, we performed a reciprocal best hit (RBH) analysis of protein BLAST searches among the genes of the 63 families (Methods). This identified 14 genes across different families that were close homologs (E-value < 10^-20^), which we reassigned accordingly (Supplementary table 2). For example, ribosomal binding factor A (*rbfA*) of *B. uniformis* was grouped in a separate COG than its orthologs in *P. merdae* and *P. vulgatus*, despite all three being essential, consistent with findings for *rbfA* in *E. coli*^25^. After RBH-based reassignment, 438 families contained at least one essential gene. Of these, 275 families included essential genes from all three species, while the remaining 163 showed variable conservation and/or essentiality across species (Fig. 3B). The essentiality calls were robust, as similar distributions of conserved and variable essential genes where observed when genes were grouped by their PFAM association (Extended Data Fig. 4A).

The majority of essential genes are conserved across the three species and including essential genes previously identified in *B. thetaiotaomicron*^21^ preserved 196 of the 275 (71%) essential gene families (Extended Data. Fig. 4B). Hence, these core essential genes are likely to be conserved across most Bacteroidales and were enriched in core cellular processes, such as translation, cell division and metabolism, but also included genes of unknown function (Fig. 3C, Extended Data. Fig. 4C, Supplementary table 1). In contrast, only 20 genes were uniquely essential in one species and lacked orthologs in the other two species according to the eggNOG COG assignment and our stringent RBH analysis. More than half of the unique essential genes have poor functional annotations (Supplementary table 1). An example of an annotated unique essential gene is an antitoxin in *B. uniformis* (BACUNI_03224), which is part of a putative RnlA/RnlB toxin-antitoxin system.

**Fig. 4:**
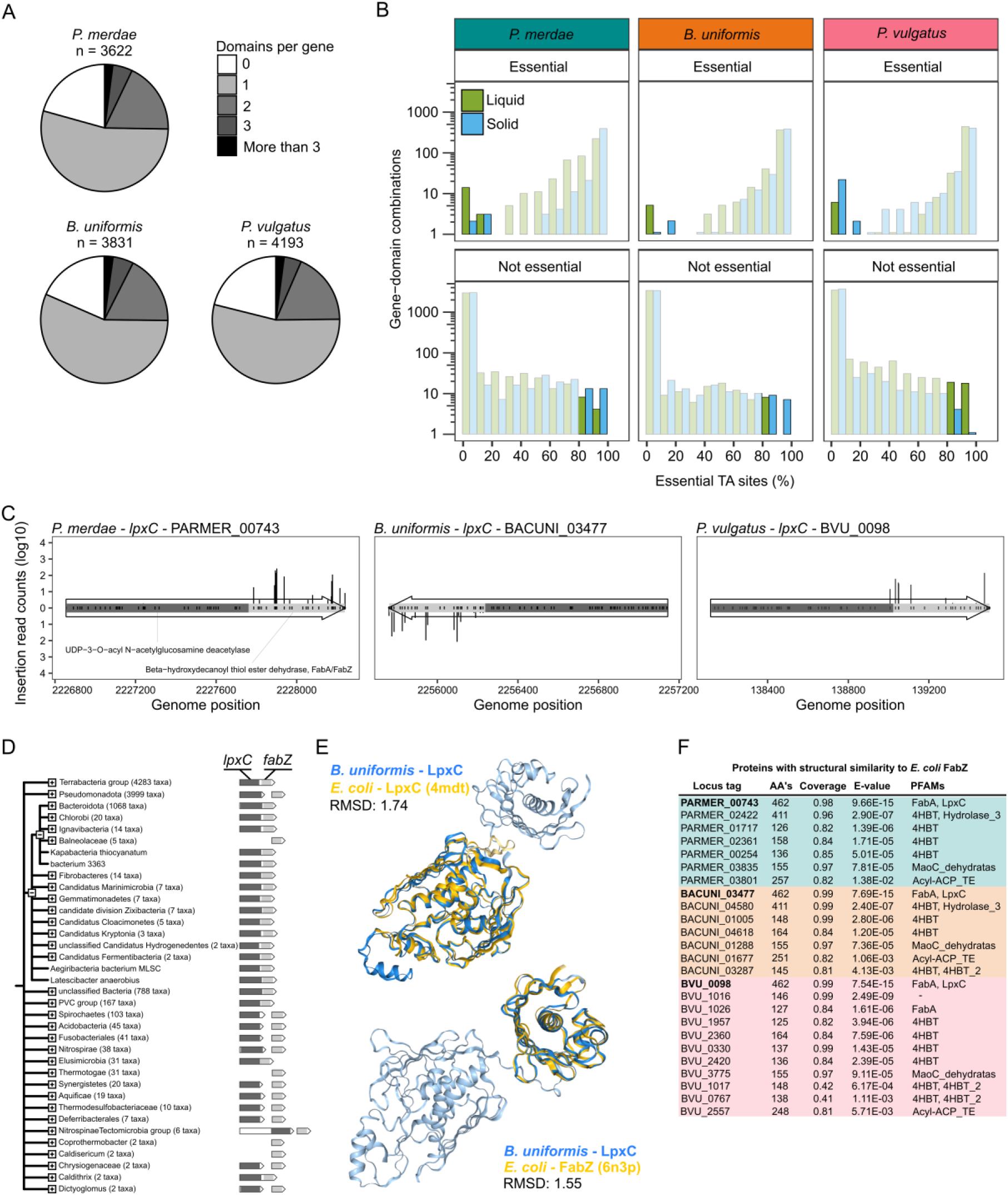
Identification of essential protein domains with transposon mutant data. **A)** Most genes of the three species encode a single protein domain. Genes are grouped by the number of their encoded protein domains per species, according to domain identification using InterProScan. *N* indicates the total number of genes analyzed. **B)** Some essential and non-essential genes encode non-essential and essential protein domains. respectively. Plotted are domains with ≥10 TA sites from genes classified as essential or non-essential, binned by the fraction of TA sites predicted to be essential using the HMM method in TRANSIT (bin width = 10%). Highlighted bars indicate cases where domains and their host gene have contrasting essentiality: non-essential domains (<20% essential sites) within essential genes (top row) and essential domains (>80% essential sites) within non-essential genes (bottom row). **C)** *lpxC* encodes two protein domains in Bacteroidales with only the one involved in lipid A biosynthesis being essential. The second domain encoding *fabZ* and involved in fatty acid biosynthesis tolerates insertions. Shown are the insertions and their read counts in the two domains of *lpxC* – domains shown in gray colors. Insertions are shown as vertical black bars. Gene arrows indicate coding direction. Transposon insertions into the forward genomic strand (+) are shown above the arrow, and insertions on the reverse strand (−) below. Black tick marks within the domain bar indicate TA sites. **D)** *lpxC* (dark gray) and *fabZ* (light gray) are fused in Bacteroidota, but appear as separate genes in different taxonomic groups of bacteria, including Pseudomonadota (formerly Proteobacteria). Taxonomic groups that show the fusion gene may include (limited) species in which the genes are separate, but the opposite does not occur. Figure is adapted from output generated by STRING-db version 12.0^110^. **E)** AlphaFold2 model of *B. uniformis* LpxC (BACUNI_03477) fusion protein (blue tones) superimposed on the crystal structures of either *E. coli* LpxC or FabZ (gold) using Foldseek. RMSD: Root Mean Square Deviation. **F)** Several proteins of *P. merdae* (PARMER), *B. uniformis* (BACUNI) and *P. vulgatus* (BVU) are predicted by Foldseek to fold similarly as *E. coli* FabZ chain A (PDB: 6n3p). Locus tags in bold are the *lpxC* genes shown in panel C. AA’s: number of amino acid residues; Coverage: fraction of protein that aligned to *E. coli* FabZ; E-value: Expect-value; PFAMs: protein families to which the protein belongs. 4HBT: 4-hydroxybenzoyl-CoA thioesterase; Acyl-ACP_TE: acyl-acyl carrier protein (ACP) thioesterases (TE).

Interestingly, the 163 gene families with non-conserved essentiality were enriched in coenzyme transport and metabolism (Fig. 3C), suggesting that essentiality can be bypassed either through redundancy (isoenzymes) or via alternative metabolic routes. An example of the latter involves *nadA*, *nadB* and *nadC*, which were assigned as essential in *P. merdae* and *P. vulgatus*, but not in *B. uniformis* and in *B. thetaiotaomicron* according to a prior study^21^ (Fig. 3D). These genes encode enzymes that synthesize nicotinate mononucleotide (NaMN) from aspartate^57^, a key precursor in the biosynthesis of the essential cofactors NAD and NADP. Loss-of-function mutations in *nadA-C* cause niacin auxotrophy^58^. However, some bacteria can bypass this requirement through a salvage pathway, in which NaMN is produced from exogenous nicotinic acid, likely present in complex mGAM media, via the enzyme PncB^59^. Indeed, *pcnB* is absent from *P. merdae* and *P. vulgatus*, explaining the essentiality of *nadA– C*, but it is present in both *B. uniformis* and *B. thetaiotaomicron*, rendering the route involving *nadA–C* dispensable (Fig. 3D). To investigate further whether functional redundancy contributes to differences in gene essentiality across species in general, we compared the number of genes per species in conserved versus non-conserved essential gene families. Paralogs resulting from gene duplications are known to buffer against gene essentiality^60^. Consistently, conserved essential families contained fewer genes per species than families in which essentiality was not conserved, suggesting that the differences in number of paralogs may explain several cases of non-conserved essentiality across the three species (Fig. 3E). An example is a putative UDP-glucose-6-dehydrogenase in family 2FMZ9 that is essential in *P. vulgatus*, which has only a single copy of this gene, but is dispensable in *P. merdae* and *B. uniformis*, which have two and three paralogs respectively (Extended Data. Fig. 4D).

In summary, the essentialome in the three Bacteroidales genera is strongly conserved and comprised 825 essential genes from 275 families, amounting to ∼80% of all essential genes of the three species combined. Species-specific differences in essentiality primarily reflected functional redundancy through gene duplication and alternative metabolic strategies.

### Genes encoding essential protein domains

Having highly saturated transposon libraries we next determined essentiality at the sub-gene level of encoded protein domains, as done previously for *E. coli*^25^ and *C. crescentus*^24^. To assign domains, we complemented gene annotations with protein domain predictions obtained from InterProScan^61^. More than half of all genes encoded a single protein domain, while about 15–20% encoded either no domain or two domains in the three species – very few proteins had more than two domains (Fig. 4A). Since several genes lacked any InterPro-annotated domain or their domains covered only a fraction of the full-length protein, we classified unannotated sections and inter-domain sequences as independent “regions” for analysis. We used the HMM method of TRANSIT to predict the essentiality state of each TA site in the updated domain-and-region annotated genome. As expected, the majority of TA sites in domains encoded by essential genes were essential, whereas TA sites in non-essential genes were predominantly marked as non-essential (Fig. 4B). However, in some cases, essential domains or regions (defined as having >80% essential TA sites) were embedded within non-essential genes, ranging from 5 to 31 instances in the *P. vulgatus* solid and liquid libraries, respectively. Conversely, we also identified non-essential domains or regions (<20% essential TA sites) within essential genes, ranging from 2 cases in the *B. uniformis* solid library to 20 in the *P. vulgatus* solid library (Supplementary table 3).

We specifically focused on a subset of genes with more than one protein domain/region (∼25% of all genes in each species; Fig. 4A) that contained both essential and non-essential domains/regions predicted with high confidence (Methods). To increase robustness, we required each domain/region to contain at least 10 TA sites, and that domain essentiality calls agreed between the liquid and solid libraries. In this way we identified 30 genes as such ‘domain-specific essentials’ (8 in *P. merdae*, 8 in *B. uniformis,* and 14 in *P. vulgatus*), of which 3 overlapped between at least 2 species (Supplementary table 3). Although some of these may be false positives due to sporadic insertions at the termini of an essential gene (see also Discussion), most likely represent genuine examples of protein domain essentiality.

One such example is DNA polymerase I (*polA*), which exhibited a similar insertion pattern across the three species: the N-terminal 5′→3′ exonuclease domain was essential, while the proofreading (3′→5′ exonuclease) and polymerase domains tolerated insertions (Extended Data Fig. 5A). This finding recapitulates known biology of PolA, as the 5’→3’ exonuclease activity is vital for DNA replication by processing RNA primers during Okazaki fragment maturation^62^. In contrast, proofreading and DNA polymerase activities can be also performed by other polymerases (e.g., Pol III and Pol II), and hence are dispensable.

Another example of domain-specific essentiality comes from an uncharacterized gene family, uniquely present in the Bacteroidota phylum. Members of this family (BACUNI_00679, BVU_3062, PARMER_04194) were identified as domain-specific essential genes in all three species (Extended Data Fig. 5B-C). These genes encode an N-terminal lipoprotein signal peptide, a central domain of unknown function (DUF4296) and a C-terminal small beta-barrel-like domain with a disordered region as predicted by Alphafold^63^ (Extended Data Fig. 5D). Essentiality mapping revealed that the DUF4296 is indispensable, whereas the C-terminal portion of the protein tolerated insertions. In all three species, this gene is located downstream of *lspA* (Extended Data Fig. 5C), which encodes an essential lipoprotein signal peptidase. The preserved genomic context and the matched essentiality suggest that *lspA* and the uncharacterized protein, may be functionally related.

A third example of a domain-specific essential gene is *lpxC*, which was annotated as encoding a multifunctional fusion protein. This gene harbored both essential and non-essential domains in *P. merdae* with a similar domain-specific transposon insertion pattern in *B. uniformis* and *P. vulgatus* (Fig. 4C). In the Bacteroidota and many other phyla, this gene appears to be a fusion of *lpxC* (N-terminal, predicted UDP-3-O-acyl-N-acetylglucosamine deacetylase activity involved in lipid A biosynthesis), and *fabZ* (C-terminal, predicted beta-hydroxyacyl-(acyl-carrier-protein) dehydratase activity involved in fatty acid biosynthesis) (Fig. 4D). The structural predictions of these domains match very well with their *E. coli* homologues (Fig. 4E, Extended Data. Fig. 6). In several other phyla, including the Pseudomonadota (formerly Proteobacteria), where most of our knowledge on the function of these genes originates, *lpxC* and *fabZ* occur as separate, distantly located genes (Fig. 4D). Both genes are essential in *E. coli*^25^ and many other bacteria, and are considered antibiotic targets^64–67^. Interestingly, in *E. coli*, pharmacological inhibition of LpxC, leads to drug resistance through mutations in FabZ^68,69^ and the two proteins have direct functional links^70^. Hence the fusion of LpxC and FabZ in Bacteroidota and other taxa may facilitate direct functional coordination between their activities. The transposon insertions tolerated in the FabZ domain suggest redundancy in the function of this domain that may be compensated by other proteins. To test this hypothesis, we searched for structural homologues of *E. coli* LpxC (PDB: 4mdt^71^) and FabZ (PDB: 6n3p^72^) in the proteomes of *B. uniformis*, *P. merdae* and *P. vulgatus*, using FoldSeek^73^. For LpxC, the fusion protein was the only high-confidence hit. In contrast, FabZ searches retrieved several proteins predicted to be structurally similar, though with lower confidence than the FabZ domain of the fusion protein (Fig. 4F & Extended Data. Fig. 6). The retrieved structural homologs are part of PFAM families, some of which are involved in other steps of fatty acid biosynthesis (e.g., BVU_1026 encodes a FabA-domain protein). One or more of these genes may compensate for loss of FabZ function in the *lpxC–fabZ* fusion transposon mutants.

In summary, our highly saturated transposon libraries reveal that essentiality is not always uniform across a protein, highlighting conserved cases of domain-specific essentiality, and pinpointing essential domains of unknown function, for future targeted functional investigation.

### Essentiality of non-coding regions

As transposons also insert in non-coding regions of the genome, we used the TRANSIT-based essentiality calling also for non-protein-coding elements in the three genomes. These elements include transfer RNAs (tRNAs) and other non-coding RNAs (ncRNAs), terminal inverted repeat elements (TIREs) of mobile elements, and intergenic regions, defined as any sequence located between annotated features (gene, tRNA, ncRNA, or TIRE). Many of these non-coding elements are relatively short, leading to unclear essentiality predictions; however, for a substantial fraction, we were able to assess essentiality: between ∼25% (TIREs) and above 90% (ncRNAs) across the three species (Fig. 5A, Supplementary table 1).

**Fig. 5:**
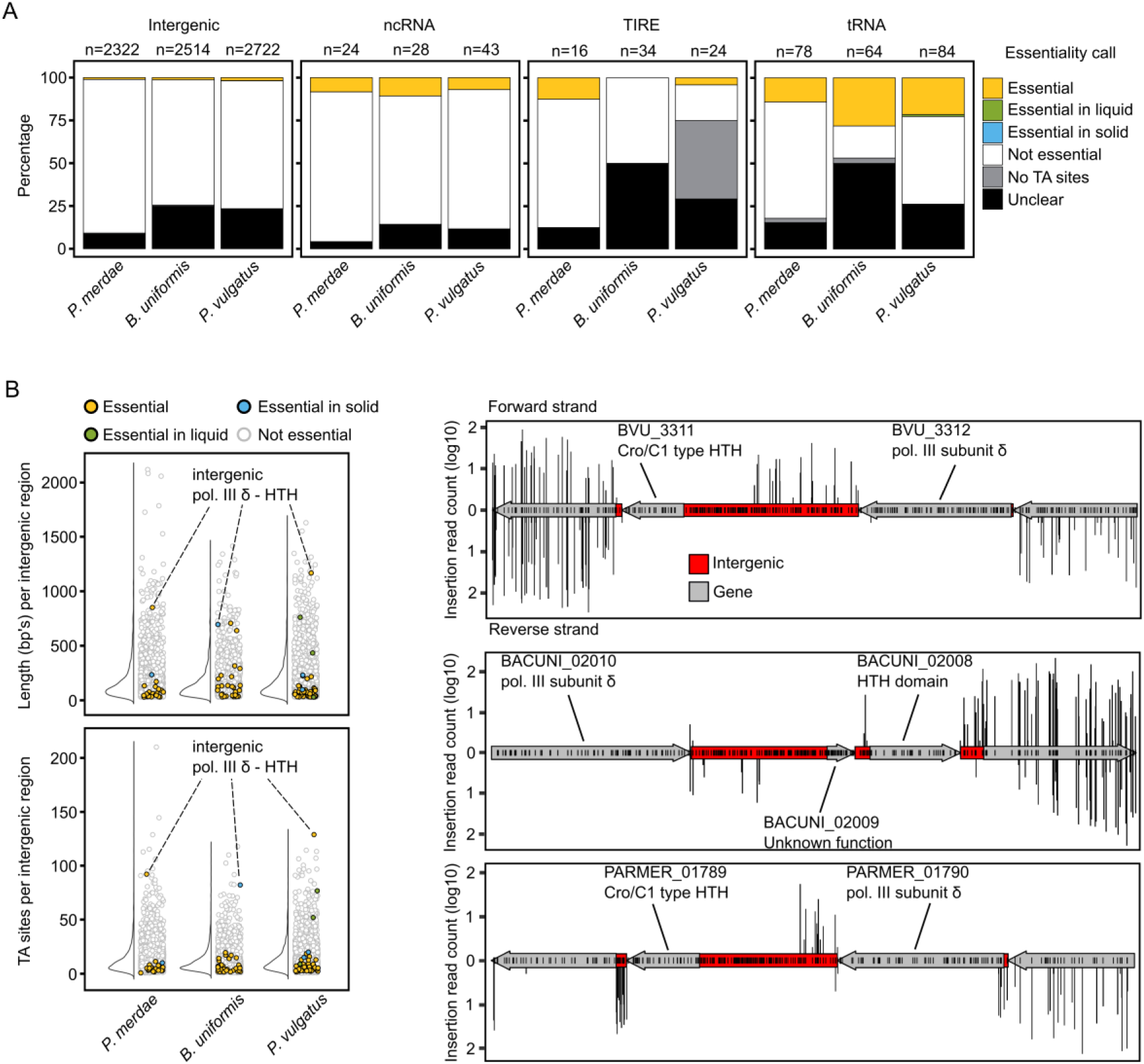
Essentiality of non-protein-coding regions. **A**) Most non-protein-coding elements are not essential. Proportion of annotated intergenic regions, non-coding RNAs (ncRNAs), terminal inverted repeat elements (TIREs), and transfer RNAs (tRNAs) with essentiality calls across the three species. **B**) A long conserved essential intergenic region between the DNA polymerase III δ subunit and a putative helix-turn-helix (HTH) transcriptional regulator. Left panels: distribution of number of TA sites in intergenic regions across the three species (function of intergenic region length), color-coded by essentiality status. Intergenic regions with no TA sites or unclear essentiality are not shown. The conserved essential intergenic region between the DNA polymerase III δ subunit and a putative helix-turn-helix (HTH) transcriptional regulator shown in the right panel is marked. Right panel: transposon insertion profiles for intergenic regions (red) and genes (gray) in *P. vulgatus* (top), *B. uniformis* (middle), and *P. merdae* (bottom). Genes and transposon insertions are depicted as in Fig. 4C.

Universally conserved non-coding elements, such as various tRNAs, the RNase P ribozyme and the signal recognition particle (SRP) RNA were also essential in the three species. Surprisingly, the transfer-messenger RNA (tmRNA) and its associated protein SmpB were essential in *B. uniformis* and *P. vulgatus*, but fully dispensable in *P. merdae*. The tmRNA– SmpB complex rescues ribosomes that are stalled by degrading the mRNA and recycling the incomplete nascent polypeptides^74^. In *E. coli*, alternative ribosome rescue pathways involving *arfA* and *arfB* allow the cells to survive in the absence of tmRNA–SmpB^75–77^. We could not find sequence or structural homologs of *arfA* or *arfB*, or duplicated copies of tmRNA or *smpB*, in any of the three species. This suggests that *P. merdae* may employ a distinct ribosome rescue mechanism that either does not exist in *B. uniformis* and *P. vulgatus*, or the same mechanism exists in all thee but can only compensate for the loss of tmRNA–SmpB complex in *P. merdae*.

We also analyzed the essentiality of intergenic regions in *P. merdae* (n=2,322), *B. uniformis* (n=2,514), and *P. vulgatus* (n=2,722). We could assess the essentiality status of 75-91% of the intergenic regions (Fig. 5A), and of those with more than five TA insertion sites (above the 25th percentile for intergenic TA site density in all three species), only 11 (*P. merdae*), 16 (*B. uniformis*), and 26 (*P. vulgatus*) were classified as essential. In most cases (11, 13, and 17 in *P. merdae*, *B. uniformis*, and *P. vulgatus*, respectively), these essential intergenic regions are located immediately upstream of essential genes (Supplementary table 1), suggesting that the transposon insertion disrupts critical regulatory sequences and interferes with essential gene expression. One notable case is an essential intergenic region conserved in all three species, located between the essential δ subunit of DNA polymerase III and a predicted essential helix-turn-helix (HTH) transcriptional regulator. This region is among the longest and most TA-rich intergenic loci (130, 83, and 93 TA sites in *P. vulgatus*, *B. uniformis* and *P. merdae* respectively) (Fig. 5B), yet we observed only 18 (*P. vulgatus*), 7 (*B. uniformis*) or 10 (*P. merdae*) insertions. No protein-coding sequence could be detected within it and thus the scarcity of insertions suggests that it may harbor an elaborate regulatory element or a functional RNA not captured by our current annotation pipeline.

These findings exemplify the potential of using dense insertion libraries for the identification of conserved and species-specific essential elements beyond protein-coding genes.

### Insertion directionality bias infers toxic modalities in genomes

Transposon libraries can also provide insights into gene interactions by studying polar effects^78,79^. We designed our transposon without a transcriptional terminator to allow for read-through transcription into downstream genes and avoid polar effects when insertions occur in the sense orientation in operons. However such a design may reduce downstream expression of operon genes in antisense insertions, where the transposon integrates opposite to a gene’s orientation, through antisense transcription (Fig. 6A). We assessed insertion directionality bias with a binomial test to identify genes showing significant enrichment of sense or antisense insertions. For each gene, we also calculated a directionality ratio, defined as the fraction of sense insertions over all (sense + antisense) insertions and normalized these ratios across libraries (Extended Data. Fig. 7A) to correct for overall insertion bias (Fig. 2D, Supplementary table 4). All libraries contained a small subset of genes with directionality ratio below 0.2 or above 0.8 (Extended Data Fig. 7A). We called significantly biased genes if they met these thresholds and had an adjusted *p*-value of < 0.01 from the binomial tests in both liquid and solid libraries. Based on these criteria we identified 114, 127 and 82 significantly biased genes in *P. vulgatus*, *B. uniformis* and *P. merdae*, respectively (Supplementary table 4).

**Fig. 6:**
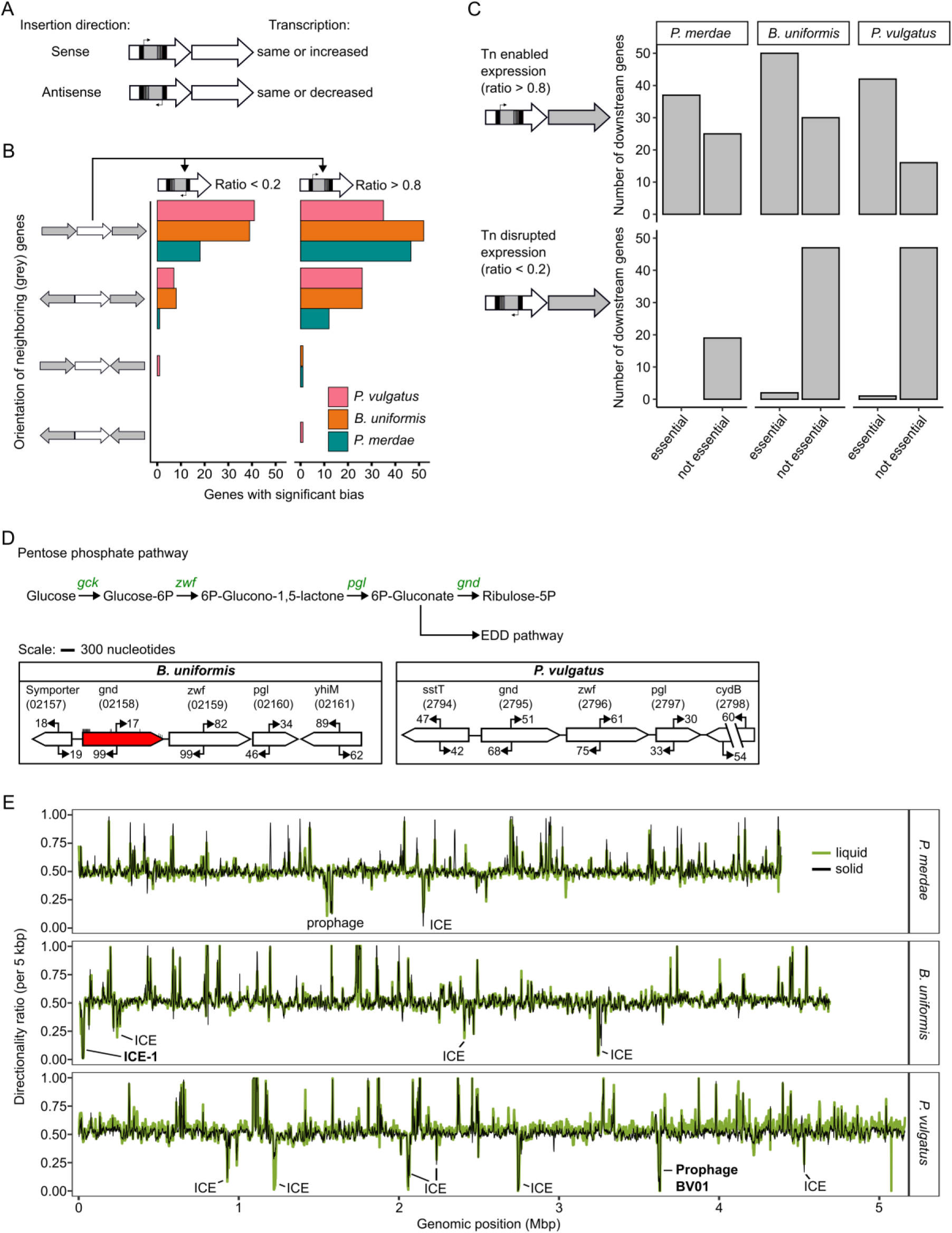
Directionality bias in transposon insertions indicates functional relations between genes. **A)** Schematic of how insertion orientation may affect downstream transcription. **B)** Genes display directionality bias only when the downstream gene has the same orientation – operon effects. Number of significantly biased genes (two-sided binomial test, Benjamini–Hochberg adjusted *p* < 0.01) grouped by orientation of their immediate neighboring genes, across the three species. The directionality ratio is defined as the fraction of sense-strand insertions over all insertions in a given gene. **C)** Essential genes counter-select antisense transposon insertions in upstream genes. Number of essential and non-essential genes downstream of significantly biased genes (binomial test, BH adjusted *p* < 0.01). **D)** Insertion bias in *gnd* suggests that production of the downstream pentose phosphate pathway genes is toxic in *B. uniformis*, but not in *P. vulgatus*. A partial pentose phosphate pathway map is shown. *gck*, glucokinase; *gnd*, 6-phosphogluconate dehydrogenase; *zwf*, glucose-6-phosphate 1-dehydrogenase; *pgl*, 6-phosphogluconolactonase; EDD, Entner– Doudoroff pathway. Arrows from the genes indicate the directionality and number of insertions summarized over the gene. *B. uniformis gnd* is highlighted in red showing a significant insertion bias (binomial test, BH adjusted *p* < 0.01, directionality ratio < 0.2). Insertions in B*. uniformis gnd* sense strand are shown as small tick marks indication mostly insertions at the edges of the gene. Genes are drawn to scale but intergenic regions are not. Locus tag identifiers (BACUNI or BVU) are indicated below gene names in parentheses. **E)** Mobile genetic elements are enriched in genes with insertion bias to avoid downstream gene expression. Genome-wide directionality ratios were calculated for coding genes in sliding windows of 5 kbps with 500 bp steps. Regions with strong bias frequently mapped to integrative and conjugative elements (ICEs) and prophages. Elements in bold are shown in detail in Extended Data Fig. 8. The line of the solid library is thinner than that of the liquid library to improve visualization.

Genes are likely biased because the transposon impacts the expression of neighboring genes. We therefore asked whether genes with significant directionality bias also showed a specific orientation of their immediate neighbors. Mapping the four possible orientations of neighboring genes revealed that nearly all biased genes are followed by a downstream gene oriented in the same direction, and hence likely part of the same operon (Fig. 6B; Supplementary table 4). This suggests that bias largely reflects whether transposon insertions enable (ratio > 0.8) or prevent (ratio < 0.2) expression of downstream genes of the same operon. Consistent with this, most of the genes that are downstream of a biased gene with ratio > 0.8 (less antisense insertions) were found to be essential (Fig. 6C). Hence antisense insertions are less well tolerated when the downstream gene is essential, by impairing its expression.

Conversely, genes with a directionality ratio < 0.2 (few sense insertions) showed a depletion of essential genes downstream (Fig. 6C). In these cases, (higher) expression of the downstream gene, while the upstream gene is disrupted by the transposon, is less well tolerated, pointing to a toxic role for the downstream gene product. Toxic modalities are known to occur in bacterial genomes, often in the form of toxin-antitoxin (TA) systems^78,80^. Across the three species, we identified 28 annotated antitoxins (Methods and Supplementary table 4) of which only 7 (2 in *P. merdae*, 3 in *B. uniformis* and 2 in *P. vulgatus*) were positioned upstream and in the same orientation as another gene, often an annotated toxin. Among these, two (PARMER_02862, BACUNI_04310) showed significant bias against downstream expression (Extended Data Fig. 7B). A third candidate antitoxin, not previously annotated as such, was identified in *B. uniformis* (BACUNI_01541) based on its bias and position upstream of an annotated toxin (Extended Data Fig. 7B). In addition, we found two further cases of similar bias in genes annotated as phage defense factors (Extended Data Fig. 7B), suggesting analogous functions to antitoxins in these defense systems.

Among the other genes showing insertional bias against expression of downstream neighbors (ratio < 0.2; few sense insertions), we identified at least five metabolic genes (2 in *B. uniformis*, 2 in *P. merdae* and 1 in *P. vulgatus*), suggesting that transposon disruption may lead to build-up of toxic metabolic intermediates^81,82^. One such example is *gnd* (6-phosphogluconate dehydrogenase) in *B. uniformis*, for which insertional bias is to potentially avoid expression of the downstream pentose phosphate pathway genes *zwf* (glucose-6-phosphate 1-dehydrogenase) and *pgl* (6-phosphogluconolactonase) (Fig. 6D). This suggests that in the absence of functional Gnd, transposon-driven expression of *zwf* (and *pgl*) may lead to a build-up of toxic 6-phosphate gluconate (6P-gluconate), possibly due to an inability of *B. uniformis* to shuttle 6P-gluconate into the Entner-Doudoroff pathway. A similar pattern was found in *P. merdae* (Supplementary table 4), but not in *P. vulgatus* (Fig. 6D). Additional examples include genes involved in inositol glycerophospholipid metabolism in *P. merdae* and *B. uniformis* (genes absent in *P. vulgatus*), and a putative esterase involved in ubiquinone biosynthesis specifically in *P. vulgatus* (Extended Data Fig. 7C). In *B. uniformis* we also found biased insertions in a gene encoding a WYL domain that is upstream of a Lin1244/Lin1753-like N-terminal domain-containing gene. Both genes are predicted to be part of a putative biosynthetic gene cluster of unknown function (Extended Data Fig. 7C).

Beyond individual genes, we observed that sets of biased genes often co-localized within specific genomic regions in each of the three species (Supplementary table 4). To capture this pattern more systematically, we determined directionality bias across the genome using a sliding window approach. This revealed regions with strong bias against downstream expression which often mapped to mobile genetic elements, such as integrative conjugative elements (ICE) and prophages (Fig. 6E). Within ICEs, several mobilization/transfer genes were significantly biased and we hypothesize that transposon-mediated dysregulation of expression of such genes can lead to cell death or possibly to excision of the ICE element (Extended Data Fig. 8A). Similarly, the prophage BV01 in *P. vulgatus*^83^ contained a group of genes towards the end of the prophage with strong insertional bias (Extended Data Fig. 8B). Although many genes in this region are of unknown function, it is plausible that dysregulation of (some of the) downstream gene(s) could lead to prophage excision.

In summary, transposon directionality bias can be used to identify functional dependencies between gene products and the processes in which they are involved, and to highlight the presence of potentially toxic modalities within genomes.

## Discussion

Bacteria in the order Bacteroidales are typically dominant members of the human gut microbiota, with several species implicated in both health and disease^39–41^, yet many remain understudied and poorly characterized experimentally. In this work, we developed tools and workflows that enable efficient construction of genome-wide transposon insertion libraries in diverse Bacteroidales species and strains. Using these resources, we generated saturated, barcoded mutant libraries in three key representative genera: *Bacteroides*, *Phocaeicola* and *Parabacteroides*. The tools and workflows presented here, include new broad-range transposon vectors with different selection markers, a dual-purpose cloning and conjugation donor strain, a convenient liquid-based protocol for generating libraries and computational scripts/tools for functional genomics analyses. Together, these innovations can be used to expedite the construction and analysis of barcoded mutant libraries across Bacteroidales species. Moreover, the three libraries established here and available to the community, can themselves serve as basis for large-scale gene-phenotype studies aimed at advancing our functional knowledge of this key group of gut bacteria.

Our dense transposon insertion libraries in *B. uniformis*, *P. merdae* and *P. vulgatus* enabled us to assess gene essentiality within and across these species. Identifying essential genes is valuable for understanding physiology and evolution^23,26,84^, refining genome-scale metabolic models^85^, predicting antibiotic sensitivities and novel targets^86^, and for engineering purposes such as utilizing auxotrophies^87,88^. Although gene essentiality has been mapped in diverse bacteria, cross-species comparisons are notoriously hard because different metrics and methods often produce false negative and positive calls^89^. To address this, we devised a systematic framework based on available bioinformatic tools, which can be used with additional mariner transposon datasets (as demonstrated here by including data from a *B. thetaiotaomicron* library^21^). Most essential gene families were involved in core processes and thus were essential across all three species, but some exhibited species-specific essentiality. We showed that species differences in essentiality often arise from functional redundancy and rerouting via alternative metabolic routes. Families with genes of unknown function were found both among conserved and species-specific essential genes.

In addition to determining gene essentiality, we mined our TnSeq dataset to systematically investigate essentiality of non-protein-coding regions, protein domains and interactions among neighboring genes using directionality bias. This revealed essential intergenic regions, critical protein domains, and indications for metabolic and mobile genetic element toxicities as a consequence of pathway disruption or mobile element activation. Such findings underscore the utility of dense transposon library TnSeq datasets for diverse functional genomics analyses. Looking ahead, applying similar analyses to new TnSeq datasets in additional species, or re-analyzing existing datasets with improved annotations for protein domains^90^, coding and non-coding genes^91^ and regulatory elements like promoters and terminators^92,93^, may yield deeper insight into the conservation and molecular basis of non-coding and protein-domain essentiality, as well as gene-dysregulation-mediated toxicity.

Our tools and approaches also have limitations. Although our transposon vector design includes promoters shown to be active across multiple species^21,43,44^, some strains did not yield transconjugants. In these cases, low conjugation efficiency is likely due to either cell envelope incompatibilities with the RP4 conjugation machinery^94,95^ or more likely due to host defense mechanisms that block incoming foreign DNA, e.g., restriction-modification enzymes.

While strain-specific strategies can sometimes overcome these barriers, broader solutions have been developed such as mimicking DNA methylation to avoid restriction-modification in the recipient^96^ or using non-RP4 based DNA delivery systems like ICE’s^97^. Mariner transposons also have inherent limitations, the most notable being their dependency on TA dinucleotide sites for insertion. This bias reduces their ability to target regions sparse in TA sites. In addition, essential genes cannot be directly studied in transposon library screens, as insertions in these regions are by definition lethal. Complementary approaches such as CRISPR interference (CRISPRi) are better suited for probing essential genes and small elements, such as ncRNAs, and have been adapted for *Bacteroides* species^98,99^.

Overall, there is a pressing need for the expansion of systematic genetic tools and genome-wide resources for gut bacteria, if we are to uncover the function and organization of genes in these non-model organisms. In this work, we provide such tools, analytical frameworks and resources for three key representatives of the human gut microbiota. Similar and complementary resources are vital for many other species and genera. This could be the basis for future large-scale genotype-phenotype mapping experiments to accelerate functional annotation and map genetic networks. As shown in other species^13,14,16,21,22^ such efforts can provide a broader knowledge base for gene function discovery and for uncovering species-specific aspects of cell physiology. Collectively such efforts will provide a foundation for deeper understanding of the gut microbiome and open new avenues for its modulation.

## Methods and materials

### Bacterial strains, growth conditions and antibiotic sensitivity testing

Bacteroidales strains used in this study are reported in Supplementary table 5. Some strains were isolated from fecal samples from healthy donors at the EMBL. The local Bioethics Internal Advisory Committee approved all experiments involving human stool-derived material. Informed consent was obtained from all donors (BIAC2015-009). All Bacteroidales strains were grown at 37°C on modified Gifu Anaerobic Media (mGAM, Nissui Pharmaceutical; 05433) in an anaerobic (12% CO_2_, 1.5% H_2_, 86.5% N_2_) vinyl chamber (Coy Laboratory Products Inc.). mGAM media (solid with 2% agar, or liquid) was placed in the anaerobic chamber to become anoxic 24 hours before use. *Escherichia coli* S17-1, *E. coli* EC100D*pir*+ (Lucigen/Epicentre) and *E. coli* DATC were grown on Lysogeny Broth (LB) at 37°C. For *E. coli* DATC, LB was supplemented with 0.3 mM diaminopimelic acid (DAP, Sigma-Aldrich; 33240). Generation of the *E. coli* DATC (available at DSMZ; 116187) conjugation donor strain is described in Bobonis *et al.* 2024^49^. To determine the sensitivity of the 32 Bacteroidales strains to erythromycin (Sigma-Aldrich; E5389), chloramphenicol (Sigma-Aldrich; C0378) and tetracycline (Sigma-Aldrich; 87128), 24-hour cultures (stationary phase) grown in mGAM were diluted in a 96-well plate 1:500 into 100 µl fresh mGAM containing antibiotics (50 µg/ml to 0.78 µg/ml by 2-fold serial decrease) and grown at 37°C. Optical density was measured every 30 minutes at 578 nm for 48 hours in an Epoch2 plate reader (Agilent) connected to a BioStack plate stacker (Agilent) installed in an anaerobic chamber.

### Construction of transposon vectors

The transposon vector pCV006 (*ermG*) was constructed using Gibson assembly of different template DNA fragments: pSAM_bt (served as vector backbone, Addgene; 112497), *cepA* hybrid promoter sequence RBF-103^44^ (synthesized, Eurofins), P_BfP1E6_ promoter sequence (taken from pWW3864^43^, kindly provided by Justin Sonnenburg) and a barcode entry sequence containing two BsmBI recognition sequences (synthesized, Eurofins). Fragments were assembled using NEBuilder HiFi DNA Assembly 2x Mastermix (New England Biolabs (NEB); E2621). The Himar1 C9 transposase gene in pSAM_bt contained a BsmBI recognition sequence, which was removed by making a synonymous nucleotide change (G to A) at position 546 relative to the starting nucleotide of the gene reading frame (A of ATG). After completing plasmid pCV006 we used NEBuilder HiFi DNA Assembly 2x Mastermix to replace the *ermG* gene for *catP* (taken from pGT-Ah7, Addgene; 122574) or *tetQ* (taken from pGT-Ah9, Addgene; 122576)^48^ to generate plasmids pCV019 and pCV020, respectively. Plasmids pCV006, pCV019 and pCV020 were verified by whole plasmid sequencing (Eurofins) and are available through Addgene (IDs as follows: pCV006: 232824; pCV019: 232825; pCV020: 232826). Annotated plasmid sequence of pCV006 is reported in Supplementary file 1.

### Barcoding of transposon vectors

One µg of pCV006 was digested with 100 units of Esp3I/BsmBI (Thermo Fisher; ER0451) in Tango buffer (Thermo Fisher Scientific (TFS); BY5) supplemented with freshly prepared DTT (Sigma-Aldrich; D0632, 1 mM final concentration) for 16 hours at 37°C, followed by 20 minutes at 65°C. Linearized plasmid was extracted from 1% agarose gel using the GeneJET Gel Extraction and DNA Cleanup Micro Kit (TFS; K0831). Single stranded DNA oligos containing a four nucleotide index in front of a 25 random nucleotide barcode were purchased from Sigma-Aldrich (cartridge purification) (Supplementary table 6). The single stranded oligo was made double stranded by PCR using a reverse primer in a reaction mixture containing: 100 pmol barcoded oligo, 500 pmol primer and 100 µl Q5® High-Fidelity 2X Master Mix (NEB; M0492). PCR program consisted of: 3 minutes at 98°C, 25 cycles of 12 seconds/cycle going from 55°C to 50°C (−0.2°C/cycle) followed by 7 minutes at 72°C. PCR reaction was purified using GeneJET Gel Extraction and DNA Cleanup Micro Kit. The linearized plasmid was barcoded and circularized through Golden Gate cloning in three identical reactions each containing: 80 ng linearized plasmid, 9 ng double stranded barcoded oligo (∼5-fold molar excess oligo:plasmid), 1 µl NEBridge® Golden Gate Assembly Kit BsmBI-v2 (NEB; E1602), 2 µl T4 ligase buffer (NEB; B0202) and water up to 20 µl total volume. The reaction mixtures were incubated for 3 hours at 42°C, after which the three reactions were pooled and purified using GeneJET Gel Extraction and DNA Cleanup Micro Kit. The barcoded plasmids were electroporated into *E. coli* EC100D*pir*+ TransForMax cells (Biozym; 190065) and expanded by growing cells until saturation at 37°C in 200 ml LB with 100 µg/ml ampicillin (Sigma-Aldrich; A9518). The barcoded plasmid library was extracted from EC100D*pir*+ cells by midi-prep (Zymo Research; D4200) and electroporated into the *E. coli* DATC conjugation donor that was grown at 37°C until saturation in 200 ml LB with 0.3 mM DAP and 100 µg/ml ampicillin. Multiple 1 ml single-use cryostocks were preserved at −80°C until use in conjugation experiments with type strains of *B. uniformis*, *P. vulgatus* or *P. merdae*.

### Whole genome sequencing and genome annotation

Genomic DNA was extracted from Bacteroidales strains isolated at the EMBL and the type strains of *B. uniformis* ATCC 8492, *P. vulgatus* ATCC 8482 and *P. merdae* ATCC 43184 (purchased from DSMZ) using the ZymoBiomics DNA miniprep kit (Zymo Research; D4300) and measured for molecular weight size distribution using FEMTO pulse (Agilent). High molecular weight DNA was fragmented using Megaruptor 3.0 (Diagenode) to a final size of 15-20 kbps. Sequencing libraries were prepared using the SMRTBell 3.0 kit (Pacific Bioscience) as per the manufacturer’s instructions. Libraries were size-selected using AMPure PB beads to remove fragments shorter than 5 kbps. Final library yield and fragment size were assessed, and libraries were pooled equimolarly and sequenced on a PacBio Sequel IIe instrument with 30 hours movie time. Genomes were assembled with Flye^100^ version 2.9-b1768. We assigned taxonomic labels using the whole genome sequence and GTDB-tk^101^ version 2.1.1. Genome sequences generated in this study have been deposited at the ENA under project number ERP180871. The *P. vulgatus* ATCC 8482 and *P. merdae* ATCC 43184 genomes were assembled into single contigs of 5.16 and 4.38 Mbps, respectively. The *B. uniformis* ATCC 8492 genome assembled into a chromosome of 4.68 Mbps and an extra chromosomal element of 22.7 kbps. Genomes were annotated using the mettannotator pipeline of the EMBL-EBI^51^, which utilizes multiple bioinformatics tools to provide information on various identified genetic elements (Supplementary table 1).

### Conjugation of transposon vectors to Bacteroidales strains

Bacteroidales recipients and *E. coli* DATC donor carrying pCV006, pCV019 or pCV020 were grown at 37°C from a single colony anaerobically in mGAM (Bacteroidales) or aerobically in LB (DATC, supplemented with 0.3 mM DAP and 100 µg/ml ampicillin) for 16-20 hours. Recipient and donor cultures were diluted 100-fold and 250-fold, respectively, in fresh media and grown for 4 hours in same conditions as overnight cultures. Donor cells were washed twice with LB without ampicillin by centrifuging at 3225 *g* for 5 min at RT. Eight OD600nm units of recipient and one OD unit of donor were collected by centrifuging cultures at 3225 *g* for 5 minutes. Pellets were resuspended in 25 µl mGAM, and recipient and donor cells were mixed and placed as 50 µl spots on mGAM agar with 0.3 mM DAP. Cells were left to conjugate for 16-20 hours at 37°C under aerobic conditions. Cell mixtures were scraped from agar into 1 ml mGAM and washed twice with mGAM without DAP by centrifugation at 3225 *g* for 5 minutes at room temperature. Pellets were resuspended in 1 ml anoxic mGAM without DAP and left to recover for 1 hour at 37°C in an anaerobic chamber. Recovered cultures were diluted 50-fold and 500-fold in mGAM, and 100 µl was plated on anoxic mGAM agar containing 25 µg/ml erythromycin, chloramphenicol or tetracycline. Forty-eight hours after incubation at 37°C in an anaerobic chamber, colonies were counted.

### Construction of full-scale mutant libraries with outgrowth in solid or liquid media

Saturated *B. uniformis* ATCC 8492 and *P. vulgatus* ATCC 8492 cultures grown anaerobically at 37°C for 24 hours from a single colony were diluted 1:100 in 10 ml MGAM and grown for 4 hours. Three cryovials of *E. coli* DATC carrying barcoded pCV006 were thawed and inoculated in 50 ml LB (supplemented with 0.3 mM DAP and 100 µg/ml ampicillin) and grown for 4 hours at 37°C in aerobic conditions with shaking at 180 rpm. Donor cells were washed twice with LB without ampicillin by centrifuging at 3225 *g* for 5 min. Eight OD600nm units of recipient and one OD unit of donor were collected by centrifuging cultures at 3225 *g* for 5 min at RT. Pellets were resuspended in 100 µl mGAM, and recipient and donor cells were mixed in quadruplets by adding 25 µl donor to 25 µl recipient. Separate mixtures were placed as 50 µl spots on mGAM agar with 0.3 mM DAP. Cells were left to conjugate for 16 hours at 37°C under aerobic conditions. Cell mixtures were scraped from mGAM agar into 4 x 1 ml mGAM and washed twice with mGAM without DAP by centrifugation at 3225 *g* for 2 min at RT. Pellets were resuspended in 4 x 1 ml anoxic mGAM without DAP and left to recover for 1 hour at 37°C in an anaerobic chamber. The four separate cell mixtures were pooled and added to 46 ml mGAM. One ml of this was serially diluted 10-fold and plated on mGAM agar with 25 µg/ml erythromycin and 200 µg/ml gentamicin to determine the input transconjugant CFUs. For the solid library, 200 µl of diluted cell mixture was spread with glass beads on 25 petri dishes (145 mm) containing mGAM with 25 µg/ml erythromycin and 200 µg/ml gentamicin. For the liquid library, 3.5 ml of diluted cell mixture was added to 100 ml of mGAM with 25 µg/ml erythromycin and 200 µg/ml gentamicin. Solid and liquid libraries were incubated anaerobically at 37°C for 32 hours. One ml of the outgrown liquid library was serially diluted 10-fold and plated on selective mGAM to determine the output transconjugant CFUs. For the solid library, all colonies were scraped and mixed in 131.5 ml mGAM containing 25 µg/ml erythromycin and 12% glycerol. The 100 ml liquid library was mixed with 31.5 ml of 50% glycerol. Multiple single-use 1 ml cryostocks were created and stored at −80°C. The mutant library of *P. merdae* ATCC 43184 was constructed as above with the following modifications. *E. coli* donor library was thawed from a second-generation glycerol stock, after previous amplification of the original *E. coli* donor library. The time of aerobic conjugation was 14 hours. For the solid library, the conjugation mix was plated with disposable L shape spreaders in 20 x 145 mm plates and 4 x 500 mm plates, and colonies were obtained after 48 h in anaerobic culturing at 37°C. Plates were scraped and added to mGAM broth before mixing with glycerol to 24% w/v and stored as 1 ml single-use pooled *P. merdae* library aliquots.

### Library preparation for TnSeq

For *B. uniformis* and *P. vulgatus* mutant libraries, genomic DNA was extracted from one 1 ml cryovial using the ZymoBiomics DNA miniprep kit (Zymo Research; D4300), while *P. merdae* DNA was extracted with the DNAeasy PowerBiofilm kit (Qiagen, 24000-50). A two-step PCR was performed to amplify transposon-genome junctions and prepare library DNA for Illumina sequencing. Five identical PCR-1 reaction mixtures were prepared per library containing: 10 µl KAPA HiFi HotStart Readymix (Roche; KK2601), 160 nM PCR-1 forward primer, 160 nM PCR-1 reverse primer 1, 160 nM PCR-1 reverse primer 2, 200 ng of library DNA and water up to 20 µl total (see Supplementary table 6 for primer sequences). PCR-1 program was: 98°C for 5 min, 6 cycles with [98°C for 30 sec, 42°C for 30 sec and −1°C per cycle, 72°C for 90 sec], 25 cycles with [98°C for 30 sec, 45°C for 30 sec, 72°C for 90 sec], 72°C for 10 min. PCR-1 reactions were pooled, purified using the GeneJET DNA Cleanup kit (TFS; K0831) and quantified using a Qubit fluorometer (TFS). Four identical PCR-2 reaction mixtures were prepared per library containing: 25 µl KAPA HiFi HotStart Readymix, 25 nM PCR-2 forward primer 1, 25 nM PCR-2 forward primer 2, 25 nM PCR-2 forward primer 3, 25 nM PCR-2 forward primer 4, 100 nM PCR-2 reverse primer, 40 ng DNA from PCR-1, water up to 50 µl. PCR-2 program was: 98°C for 5 min, 30 cycles with [98°C for 30 sec, 52°C for 30 sec, 72°C for 90 sec], 72°C for 10 min. PCR-2 reactions were pooled, purified using the GeneJET DNA Cleanup kit (TFS; K0831) and run on a 1.5% agarose gel. The expected DNA smear was cut between 300 and 900 bps and extracted from gel using the GeneJET Gel Extraction and DNA Cleanup Micro Kit. Library quality was assessed with a Bioanalyzer on a high sensitivity DNA chip (Agilent) or Tapestation D5000 HS (Agilent). Libraries were sequenced on an Illumina NextSeq 500 system with a Mid output flow cell for 150 basepair single-end reads and 10% PhiX spike-in. Yield was at least 20 million reads per library.

### Insertion mapping using TnSeeker and barcode filtering

Transposon insertions were mapped onto the genome using TnSeeker version 1.0.6.5 (https://github.com/afombravo/tnseeker) and Bowtie2 version 2.4.4^102^ and a genome feature file (GFF) which was obtained by running the mettannotator annotation pipeline^51^. The following TnSeeker input parameters were used: --tn TACGAAGACCGGGGACTTATCATCCAACCTG (Himar-1 inverted repeat), --m 6, --b, --b1 ATGTCCACGAGGTGTAACTG (*B. uniformis* and *P. merdae*) or ATGTCCACGAGGTGTATGCA (*P. vulgatus*), --b2 CAGAATTGGGAGTCTACGAA, --b1m 3, --b2m 3, --b1p 1, --b2p 1, --ph 10, --mq 1, --ne. The TnSeeker output file “annotated_barcodes.csv” was used to determine the number of unique barcodes per gene based on rules similar to those described in Price *et al.,*^14^. Briefly, good barcodes are defined as either uniquely mapping to one genome position with 5 or more reads, or mapping to more than one location with 10 or more reads of which 3/4 or more of the reads map to one primary location and 1/8 or less of the reads map to any other location. To filter out likely erroneous barcodes generated from sequencing errors, barcodes with a Hamming distance of 3 or less compared to the barcode with the highest read count at a given position were removed.

### Analysis of gene essentiality

An overview of the computational workflow and the required files and R scripts necessary for essential gene analysis is shown in Supplementary figure 1. For the identification of essential genes, we used the TRANSIT software package^54^. The necessary .prot_table and .wig file inputs for TRANSIT were generated with custom R scripts. Only genes encoding proteins between 10 and 5000 amino acids were included in the analysis (these thresholds are set when making the .prot_table file). We used TRANSIT version 3.3.2 through the graphical user interface. To determine essential genes with the HMM method, we used the following settings: ignore N-terminus %: 5, ignore C-terminus %: 10, normalization: nonorm, replicates: sum, and with correction for genome positional bias. The resulting output was run through the HMM post-processing script (downloaded from https://github.com/mad-lab/transit/tree/master/src/HMM_conf.py) to add confidence flags to the HMM calls^103^. The HMM outputs one of four calls for each gene: essential (ES), growth defect (GD), not essential (NE) or growth advantage (GA). To create a binary essentiality call, we combined GD genes with ES genes if the GD gene had no reads. GD genes were combined with NE genes if they had reads. GA genes were combined with NE genes. To determine essential genes with the Gumbel-binomial method^104^, we used the following settings: ignore N-terminus %: 5, ignore C-terminus %: 10, samples: 20000, burn-in: 1000, trim: 1, minimum: 1, replicate: sum. Essentiality calls from the HMM and Gumbel-binomial output were consolidated per library and these calls were further consolidated between the liquid and solid libraries using a custom R script according to the algorithm shown in Extended Data. Fig. 3A.

### Analyses of essential gene conservation and functional enrichment

To determine the conservation of gene essentiality across species, we grouped genes into families by mapping the protein-coding genes of the three species to the EggNOG 5.0 database^56^ using EggNOG-mapper^105^. Gene families were defined based on COG classification at the Bacteroidia (class) taxonomic level. PFAM annotations from EggNOG-mapper were used as an alternative grouping method. Genes annotated as essential in either liquid or solid media were classified as essential, while families containing genes with an “unclear” essentiality call were excluded (n = 71). Families lacking gene members from one or two species (n = 63) were analyzed by protein BLASTP (version 2.15.0+). Protein sequences of the genes in these families were queried against the complete proteomes of *P. merdae*, *P. vulgatus*, and *B. uniformis*. For each query, the top five subject hits were selected based on the highest bitscore (ties resolved by the lowest E-value). These subject hits were subsequently used as queries in reverse BLASTP searches against the proteome of the original query species. The best hit was defined as the alignment with the highest bitscore, with ties resolved by the lowest E-value. Reciprocal best hits (RBHs) were defined as gene pairs that were each other’s best forward and reverse hits. Hits were required to meet the following thresholds: query coverage >= 80%, subject coverage >= 80%, bitscore >= 80, and E-value <= 1e-20. RBHs involving three species were consolidated such that if two genes shared the same COG assignment, the third was reassigned to that family. In cases of three different family assignments or two-species RBHs, the family of the query species was adopted. This procedure resulted in the reassignment of 14 genes, reducing the total number of essential gene families from 452 (EggNOG only) to 438 after RBH-based correction. The COG category enrichment analysis was performed using the categories listed per gene family (eggNOG COG at Bacteroidia level) from the eggNOG-mapper output with a custom R script. The COG categories were retrieved from NCBI. Enrichment was assessed using a one-sided Fisher’s exact test. *p*-values were adjusted for multiple testing using the Benjamini–Hochberg false discovery rate (FDR) method.

### Analysis of essential protein domains

To evaluate the essentiality of protein domains encoded within genes, we annotated the genome at the domain level using InterProScan outputs derived from the mettannotator pipeline and a custom R script. Briefly: for each domain prediction tool in the InterProScan output (e.g., PFAM, NCBIfam), the longest (in nucleotides) entries labeled as “Domain” or “Family” were retained. Domains were prioritized over families, and redundant annotations (e.g., domains embedded in families or vice versa) were removed. Predictions with >50% overlap were consolidated, retaining only the longest prediction (e.g., PFAM prediction retained over a shorter NCBIfam prediction). Inter-domain regions were labeled sequentially (e.g., region_1, region_2), starting from the 5’ end of each gene. A new .prot_table file was generated using the domain-annotated genome file, retaining only domains and regions ≥10 amino acids. The .prot_table was used together with library-specific .wig files of insertion counts at TA sites for domain-essentiality assessment using TRANSIT’s HMM method. For the domain-essentiality analysis, unlike in the gene-essentiality analysis, insertions in the N- or C-terminus of proteins were not ignored. The HMM output classified TA sites within domains and regions as “essential” (ES), “growth defect” (GD), “non-essential” (NE), or “growth advantage” (GA) and to create a binary call, GD sites were converted to ES sites if they had no read counts and were converted to NE sites if they had read counts. GA sites were converted to NE sites. To summarize the percentage of essential TA sites in domains and regions per gene, those genes found to be essential in either solid or liquid media were counted as essential genes. To call hits of essential genes containing non-essential protein domains, or vice versa, we selected genes with at least two domains, each represented by at least 10 TA sites and without a low-confidence HMM flag. Among these, one domain had to have the highest HMM probability of being essential, while another had to have the highest probability other than being essential. The TRANSIT’s HMM essentiality output per domain and selection of hits is included in Supplementary table 3. AlphaFold monomer-predicted structures of proteins of interest were retrieved from UniProt using their accession codes. To find structural homologues of *E. coli* FabZ and LpxC, we first generated structural predictions of all proteins of the three species using ProstT5^106^ as implemented within the Phold package and using its default settings (version 0.1.3)^107^. These predicted structures were searched against the PDB (Aug-2024) using foldseek^73^ (version 9.427df8a), using default settings for the easy-search mode. Proteins with predicted structural similarities to FabZ (6n3p^72^) and LpxC (4mdt^71^) and E-values < 0.05 were retained. Alphafold models of the hits were retrieved from UniProt using each protein’s UniProt accession and were run in the foldseek webserver (Aug-2025) for visual inspection and final model scores (RMSD, E-value, TM-score).

### Analysis of insertion directionality bias

Directionality bias was assessed using a custom R script. For each gene, the directionality ratio was calculated as the number of insertions in the gene’s sense strand divided by the total insertions in both the sense and antisense strands. Ratios were median-normalized across all libraries. Directionality bias significance was determined using a binomial test with Benjamini-Hochberg correction. Genes with an adjusted *p*-value < 0.01 and a directionality ratio < 0.2 or > 0.8 were classified as significantly biased. Putative toxin and antitoxin genes were identified in the three species using two approaches: Searching for “toxin” or “antitoxin” in the gene description fields of the .gff output files generated by mettannotator and by using TAfinder2.0^108^ via the web interface with the following parameters: BLAST E-value: 0.01; HMMER E-value: 0.01; Maximum sequence length: 500 amino acids; Maximum distance: 150 nucleotides; Maximum overlap: 50 nucleotides. The input file was a .gbk file generated by mettannotator. Supplementary table 4 includes directionality bias ratios for all genes across libraries, significantly biased genes and their immediate neighbors, canonical toxin-antitoxin genes found using text mining and TAfinder2.0, and genes with similar directionality biases to toxin-antitoxin pairs.

## Data availability

Sequencing data of the *P. merdae* mutant libraries has been deposited in the European Nucleotide Archive (ENA) at EMBL-EBI under accession number PRJEB77289. Sequencing data of *B. uniformis* and *P. vulgatus* libraries has been deposited in the ENA under accession number PRJEB98479 and will be available after 01-July-2026. Whole genome sequences of *P. merdae* ATCC43184, *B. uniformis* ATCC8492 and *P. vulgatus* ATCC8482 and strains isolated from fecal samples in this work are deposited in the ENA under study number ERP180871 and will be available after 01-July-2026.

## Code availability

TnSeeker software, used for processing of TnSeq reads, is available at https://github.com/afombravo/tnseeker. Custom Unix and R scripts to perform functional genomics analyses of a single Mariner based TnSeq library are available at https://git.embl.org/grp-typas/transposon_toolkit. See Supplementary figure 1 for a flowchart of files, scripts and softwares used for the analyses.

## Acknowledgements

We thank Vitalina Chamberlain-Evans (MRC Toxicology Unit, University of Cambridge) for helpful conversation and guidance on DNA library QC. We thank all Typas group members for helpful discussions and in particular, Tara Bartolec for running and providing Foldseek output and Martin Garrido Rodriguez-Cordoba for assistance with Alphafold modelling. We thank Justin Sonnenburg for providing the pWW3864 plasmid. We thank the EMBL genomics core facility, and in particular Vladimir Benes, Mireia Osuna Lopez and Hilal Ozgur for their help and support with whole genome and TnSeq sequencing library preparation.

## Author contributions

Conceptualization: C.V., I.R., M.Z, K.R.P., A.T. Funding acquisition: C.V., M.Z, K.R.P., A.T. Experiments: C.V., I.R., K.M., J.B. Data analysis: C.V., I.R., N.K., A.B., L.K., V.V. Writing manuscript: C.V., I.R., A.T. with edits from all authors. Visualization: C.V., I.R. Supervision: G.Z., M.Z., K.R.P., A.T.

## Extended data with legends

**Extended data Fig. 1:**
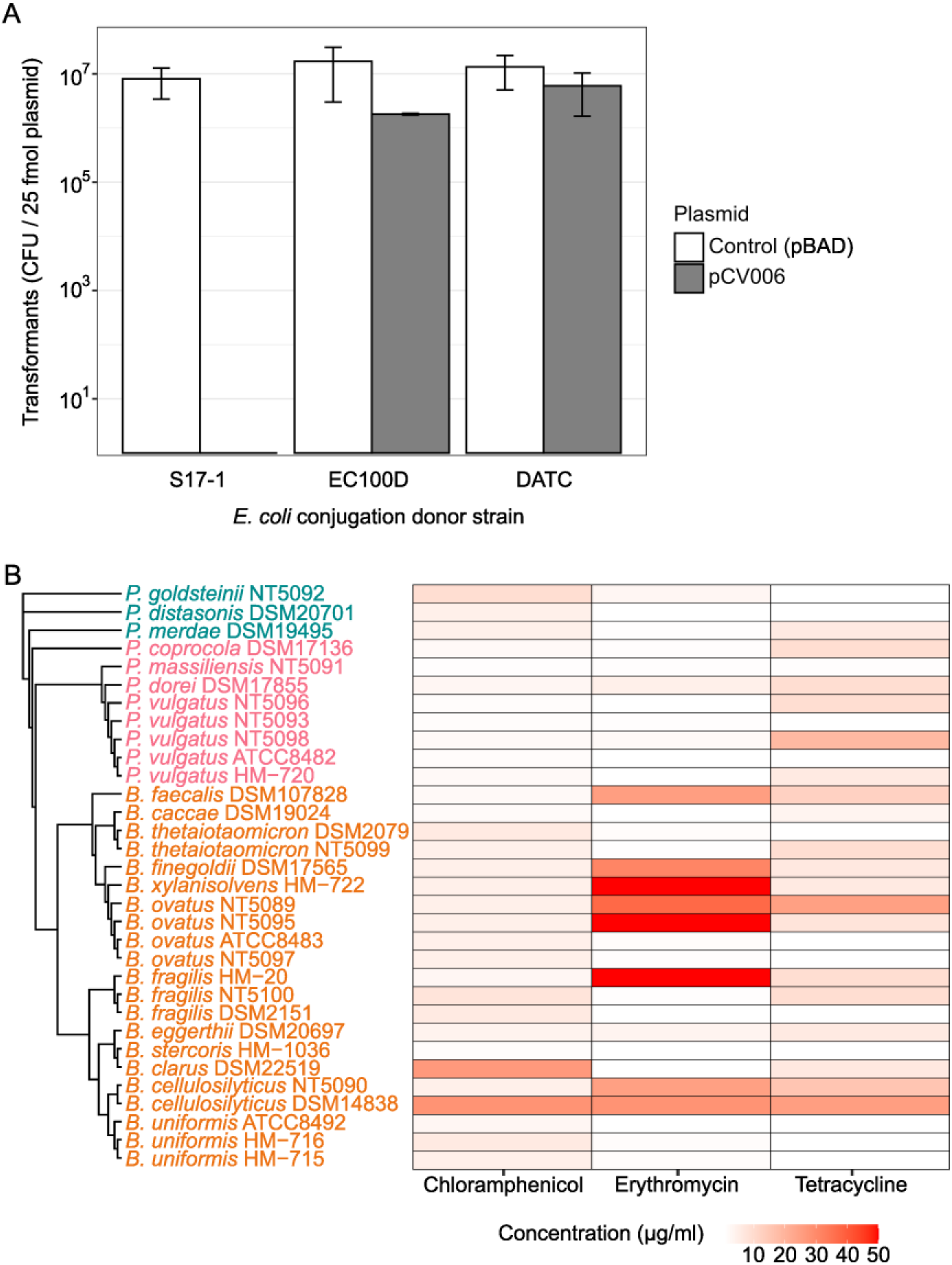
Transformation efficiency of *E. coli* DATC donor strain and antibiotic resistance profiles of Bacteroidales strains. **A)** The *E. coli* cloning-conjugation donor strain DATC can be transformed with high efficiency. Transformation efficiency of the pCV006 transposon vector or a control plasmid (pBAD) into different strains of *E. coli* – shown as CFU of ∼ 5 x 10^9^ input cells per 25 fmol plasmid. Bars indicate the mean and SEM of three biological replicates. **B)** Antibiotic resistance among strains of diverse Bacteroidales species. Shown in red shade is the highest concentration of chloramphenicol, erythromycin and tetracycline (maximum concentration tested was 50 μg/ml) for which growth was observed among 32 strains of *Parabacteroides* (green), *Phocaeicola* (pink) and *Bacteroides* (orange) species. Values indicate the mean of two biological replicates. Strains are grouped by phylogeny according to a neighbor-joining tree build on whole genome average nucleotide identity approximated with Mash version 2.3^109^.

**Extended data Fig. 2:**
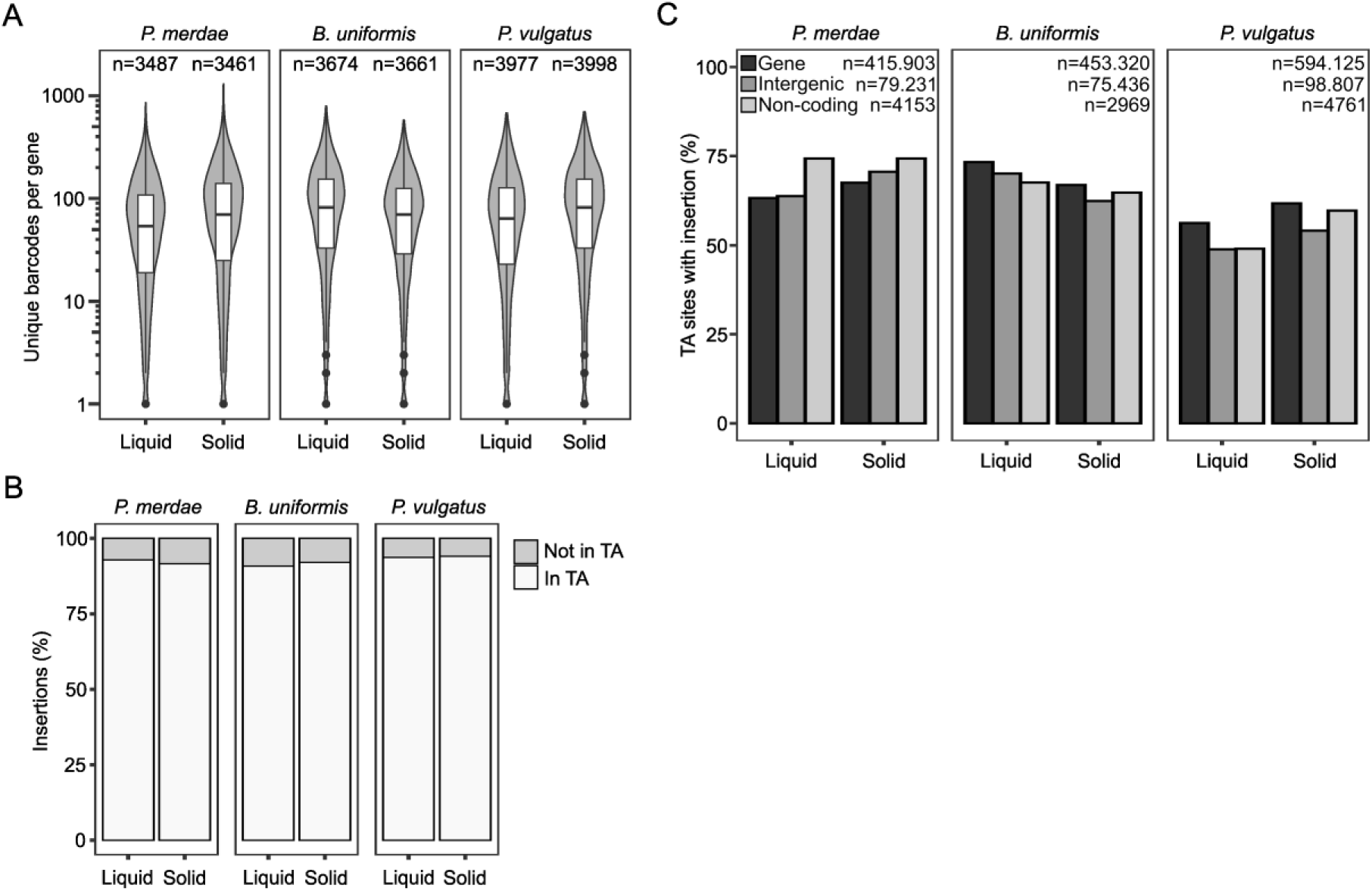
Barcode coverage and genomic distribution of insertions across transposon libraries. **A)** Almost all genes in the 6 libraries are represented by multiple unique barcodes. *N* indicates number of genes analyzed. Boxplots show the distribution of the number of unique barcodes per gene in each transposon library. The center line represents the median; box limits indicate the first and third quartiles; whiskers extend to 1.5 x the interquartile range; and points represent outliers. **B)** Percentage of insertions at TA dinucleotide versus any other site per library. **C)** Percentage of TA dinucleotide sites with insertions in genes, annotated non-coding features and intergenic regions. Inset numbers indicate the total number of TA sites per feature in each species.

**Extended data Fig. 3:**
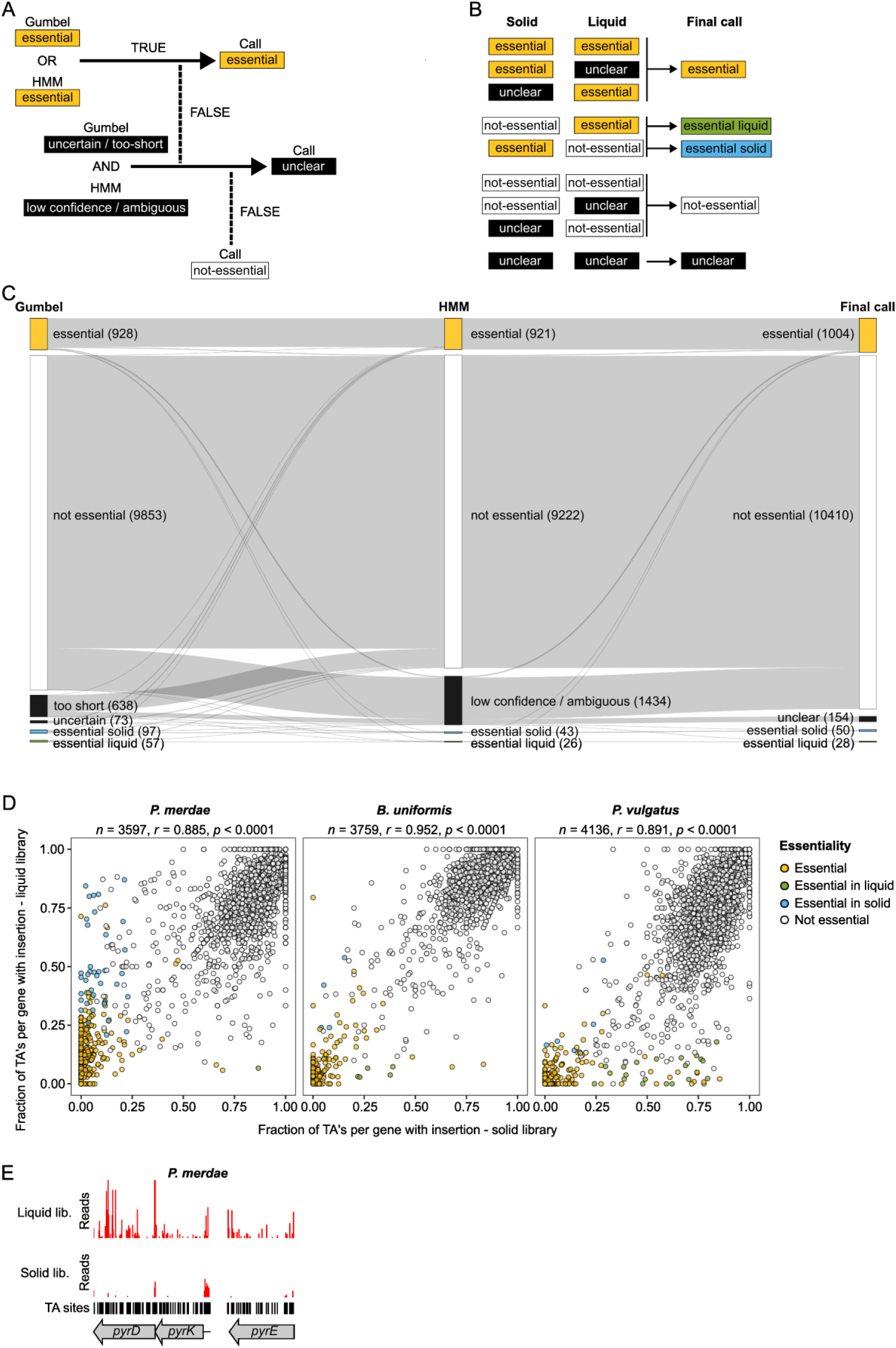
Gene essentiality calling between solid and liquid libraries. **A)** Logic used to consolidate essentiality calls from TRANSIT’s Gumbel and HMM methods post analysis. **B)** Logic used to consolidate calls from libraries grown on solid and liquid media to create the final essentiality call. **C)** Flow diagram illustrating the combined number of genes from the three species, their essentiality call by the Gumbel and HMM methods and how the consolidation between these two methods yields the final essentiality call (Supplementary table 1). **D)** Essentiality calls strongly agree between solid and liquid grown libraries. Shown are the fraction of TA dinucleotide sites per gene with transposon insertions in solid (x axis) versus liquid (y axis) libraries for the three species. *n* = number of genes, *r* = Pearson correlation coefficient, *p* = *p*-value (two-sided). **E)** Representative TRANSIT insertion plots of *P. merdae pyr* genes from pyrimidine metabolism that are essential in liquid growth conditions (in contrast to solid conditions), possibly due to complementation of auxotrophies in the pooled library.

**Extended data Fig. 4:**
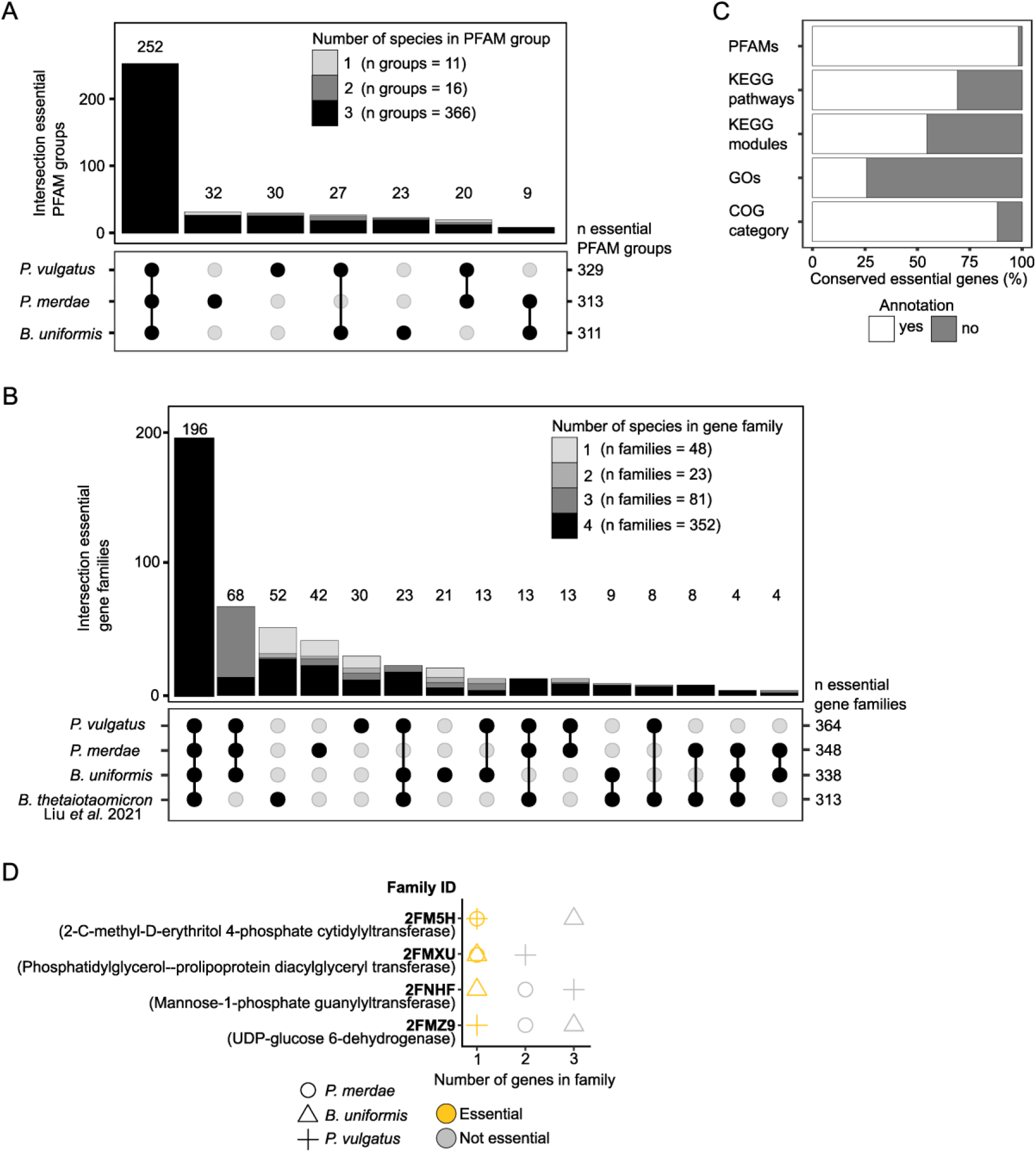
Conservation, annotation, and copy number shape essential gene families in Bacteroidales. **A)** Conservation and species-specificity of gene essentiality is preserved when genes are grouped by their annotated PFAMs instead of eggNOG COGs – plotted as in Fig. 3B. **B)** Addition of *B. thetaiotaomicron* essential genes from Liu *et al*., 2021^21^ largely preserves the core essentialome of members of the Bacteroidales. Shown are essential gene families as defined by Cluster of Orthologous Groups (COGs) at the Bacteroidia (class) taxonomic level – plotted as in Fig. 3B. **C)** Most conserved essential genes have a functional annotation, yet some lack any annotation from the indicated functional categorization resources. Shown is the percentage of genes (n = 836) that are in the 275 conserved essential gene families (some families have more than one gene per species) from Fig. 3B. **D)** Essentiality depends on gene copy number. Shown are examples of Bacteroidia taxonomy level (class) eggNOG gene families in which single-copy genes are essential, while in species for which the same family has additional members, all are not essential.

**Extended data Fig. 5:**
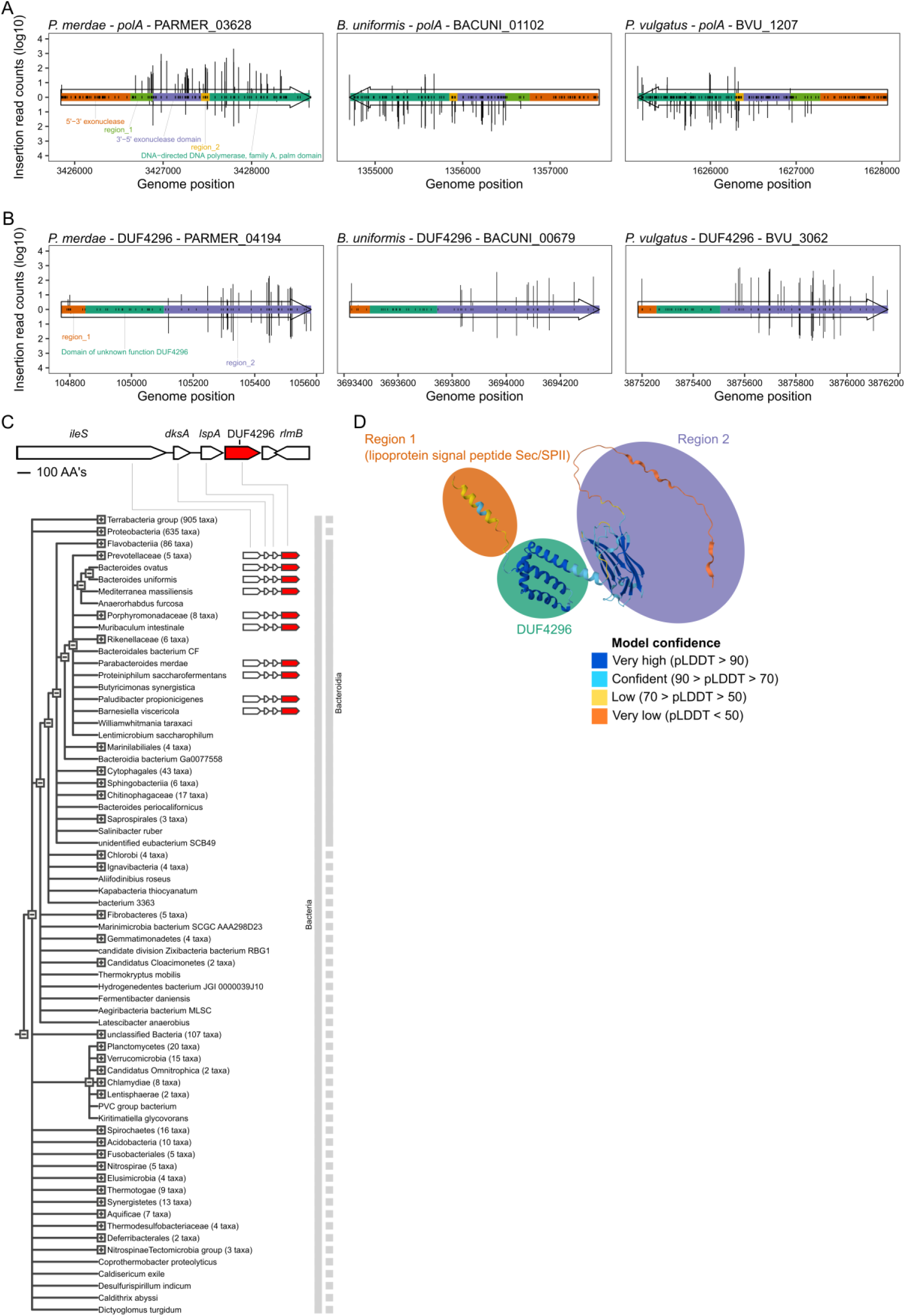
Examples of sub-gene essentiality of encoded protein domains. **A)** Only the 5’ to 3’ exonuclease domain of DNA polymerase I (*polA*) is essential. Shown are insertions and their read counts in the different domains in *polA* (color-coded and denoted in inset). Gene arrows indicate coding direction (5′→3′, sense strand). Transposon insertions into the forward genomic strand (+) are shown above the arrow, and insertions on the reverse strand (−) below. Colors within the arrow indicate the different domains and regions. Black tick marks within the arrow indicate TA sites. **B)** Insertion plot as in A of a conserved, Bacteroidota-specific uncharacterized gene with an essential domain of unknown function (DUF4296). Insertions are not tolerated within the DUF4296, but are frequent within the two adjacent regions of the gene. **C)** Presence and genomic neighbors of the uncharacterized gene with DUF4296 across different taxonomic groups of bacteria (visualization is adapted from output generated by STRING-db^110^). **D)** Predicted structure (pTM score = 0.48) of the *P. vulgatus* DUF4296 encoding gene (BVU_3062) obtained by Alphafold3^111^. Colors indicate the model confidence score pLDDT (predicted local distance difference test).

**Extended data Fig. 6:**
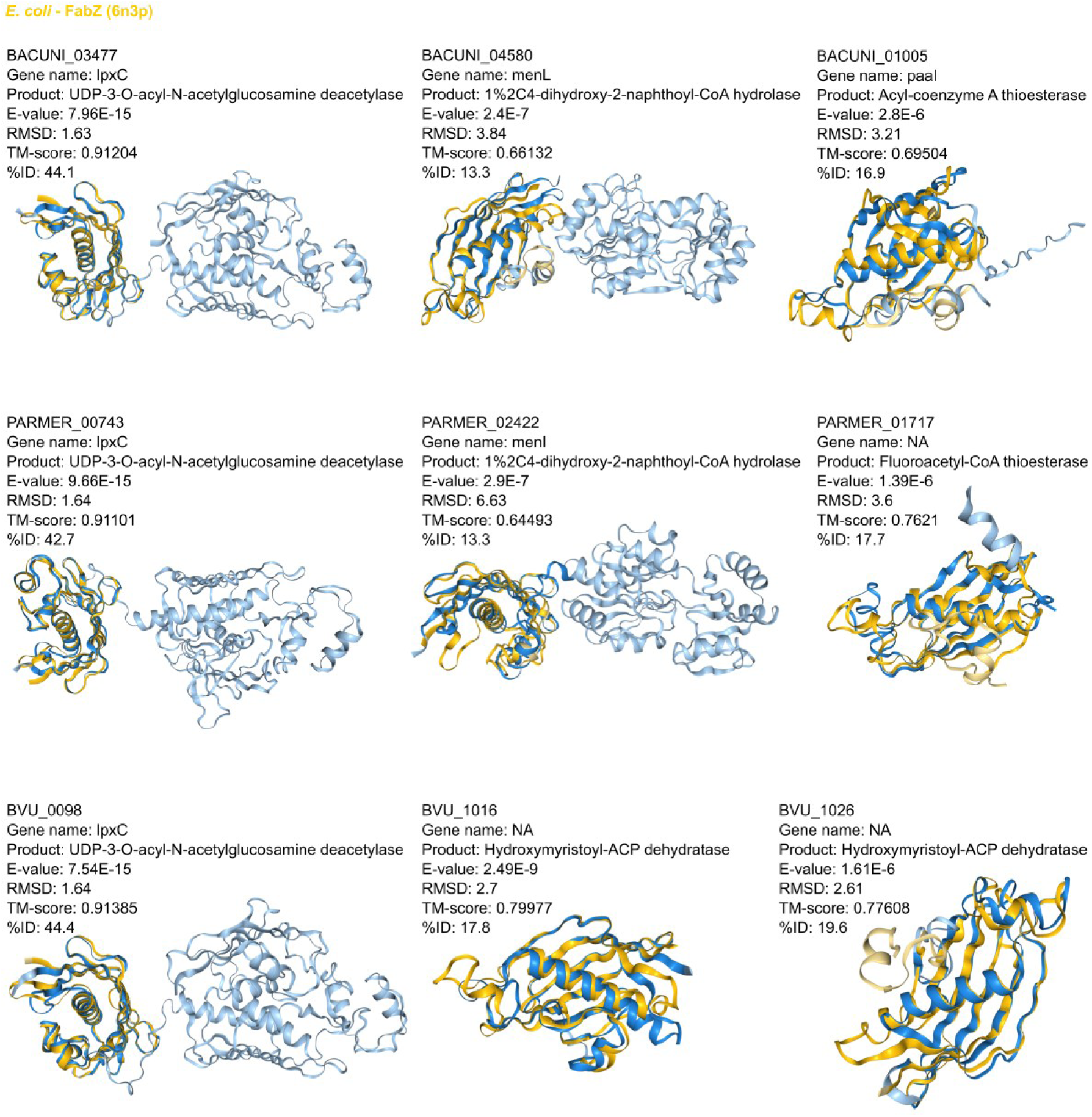
Structural similarity of Bacteroidales proteins to *E. coli* FabZ. Shown are the top three proteins of *P. merdae*, *B. uniformis* and *P. vulgatus* that are predicted by Foldseek^73^ to have the highest similarity (lowest E-value, Fig. 4F) to *E. coli* FabZ (in gold). RMSD: Root Mean Square Deviation. TM: Template Modeling score. %ID: percentage of identical amino acid residues in the alignment.

**Extended Data Fig. 7:**
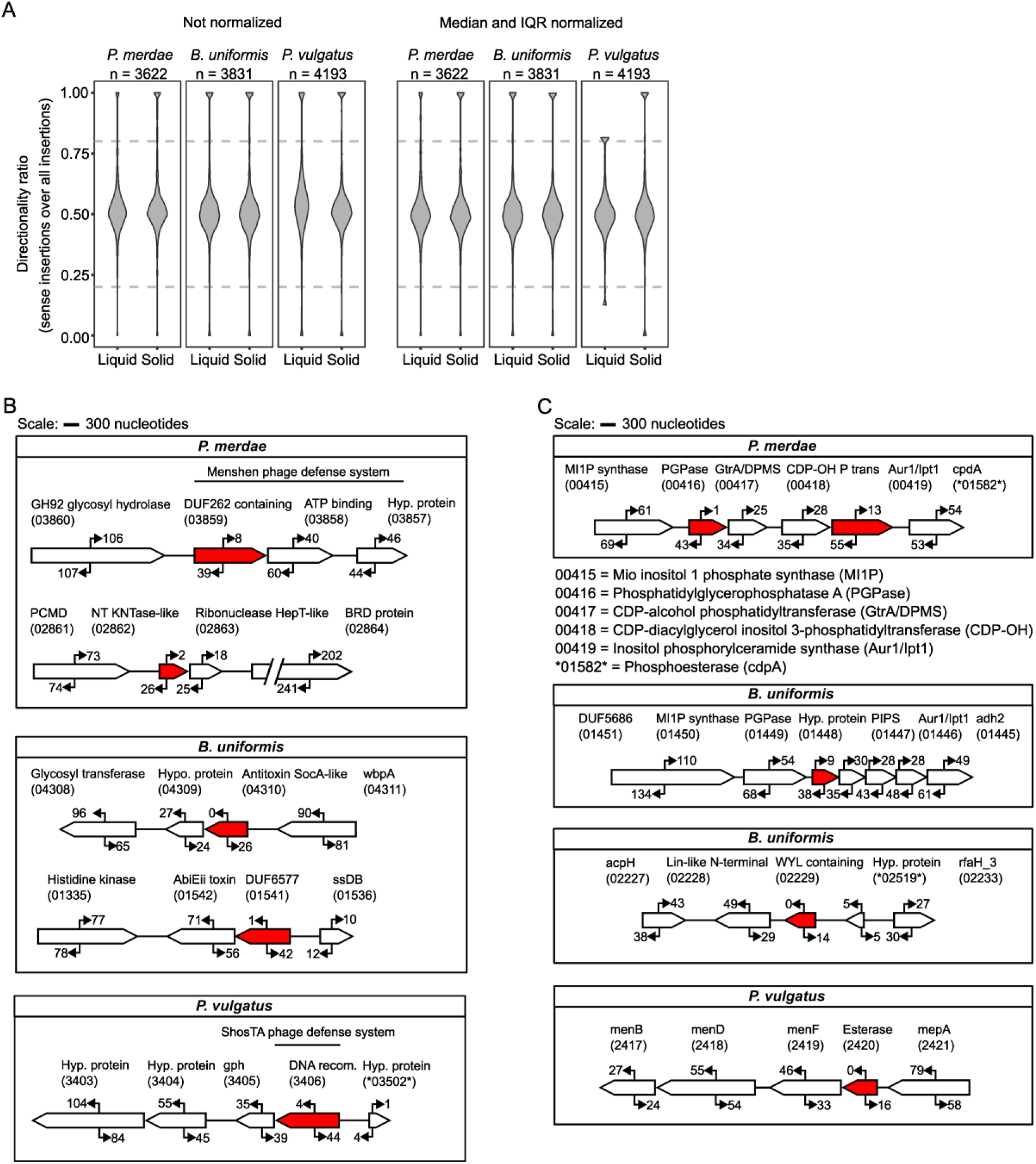
Directionality bias of transposon insertions. **A)** Some genes show strong strand bias, either favoring sense insertions (ratio > 0.8) that permit downstream expression or antisense insertions (ratio < 0.2) that suppress it. Violin plots show the distribution of directionality ratios (fraction of sense-strand insertions per gene) across all libraries, before and after median/IQR normalization. Dashed grey lines indicate 0.8 and 0.2 thresholds. **B–C)** Examples of biased genes in toxin–antitoxin and phage defense systems (B), and in metabolic pathways (C). Arrows represent the orientation and number of insertions per gene. Significantly biased genes (two-sided binomial test, Benjamini–Hochberg adjusted *p* < 0.01; directionality ratio < 0.2) are shown in red. Genes (but not intergenic regions) are drawn to scale (scale bar: 300 nt). Numbers below gene names in parentheses indicate locus tags (PARMER, BACUNI, BVU). Numbers marked with asterisks indicate genes without canonical locus tags; these instead carry PM_ATCC43184, BU_ATCC8492, or PV_ATCC8482 identifiers introduced in this study (Supplementary table 1).

**Extended Data Fig. 8:**
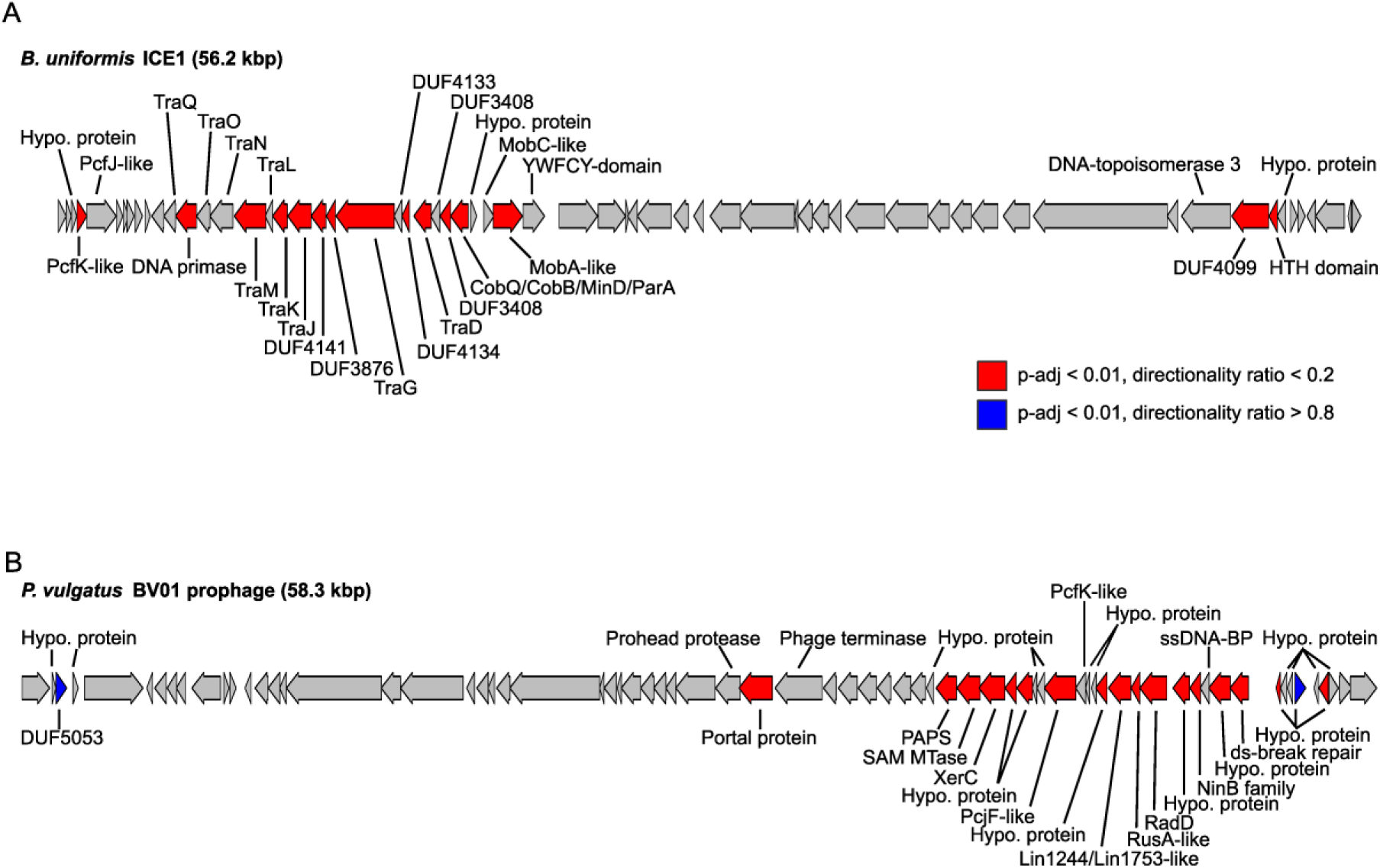
Clusters of directionality-biased genes in mobile genetic elements. **A)** An integrative and conjugative element (ICE) in *B. uniformis* contains multiple genes biased against downstream expression. **B)** The *P. vulgatus* prophage BV01 contains multiple genes near one end that are similarly biased to avoid mostly downstream gene expression. In both panels significantly biased genes are shown in red (antisense insertions; directionality ratio < 0.2) and blue (sense insertions; directionality ratio > 0.8) – significance assessed by two-sided binomial test, Benjamini–Hochberg adjusted *p* < 0.01.

## Supplementary material

**Supplementary Fig. 1:**
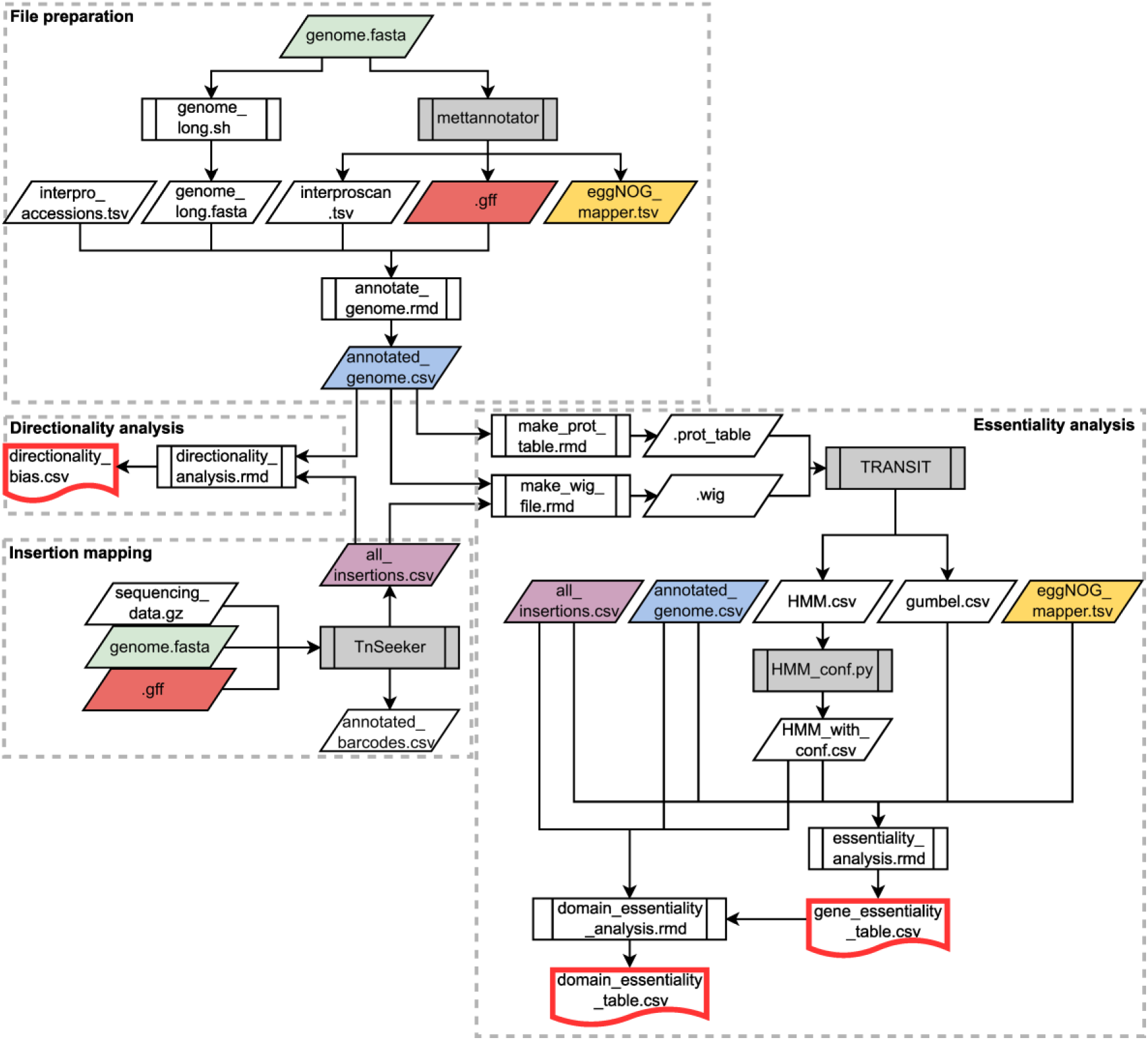
Inputs, outputs and softwares used to perform the functional genomics analyses. Flowchart depicting the input and output files and softwares and scripts for the functional genomics analyses. Recurrent files are color coded. Scripts for processes in white background are provided with this work and can be downloaded from https://git.embl.org/grp-typas/transposon_toolkit. Processes in gray background are described elsewhere, see Methods. Output files outlined in red are the final files for essentiality, domain essentiality and directionality data.

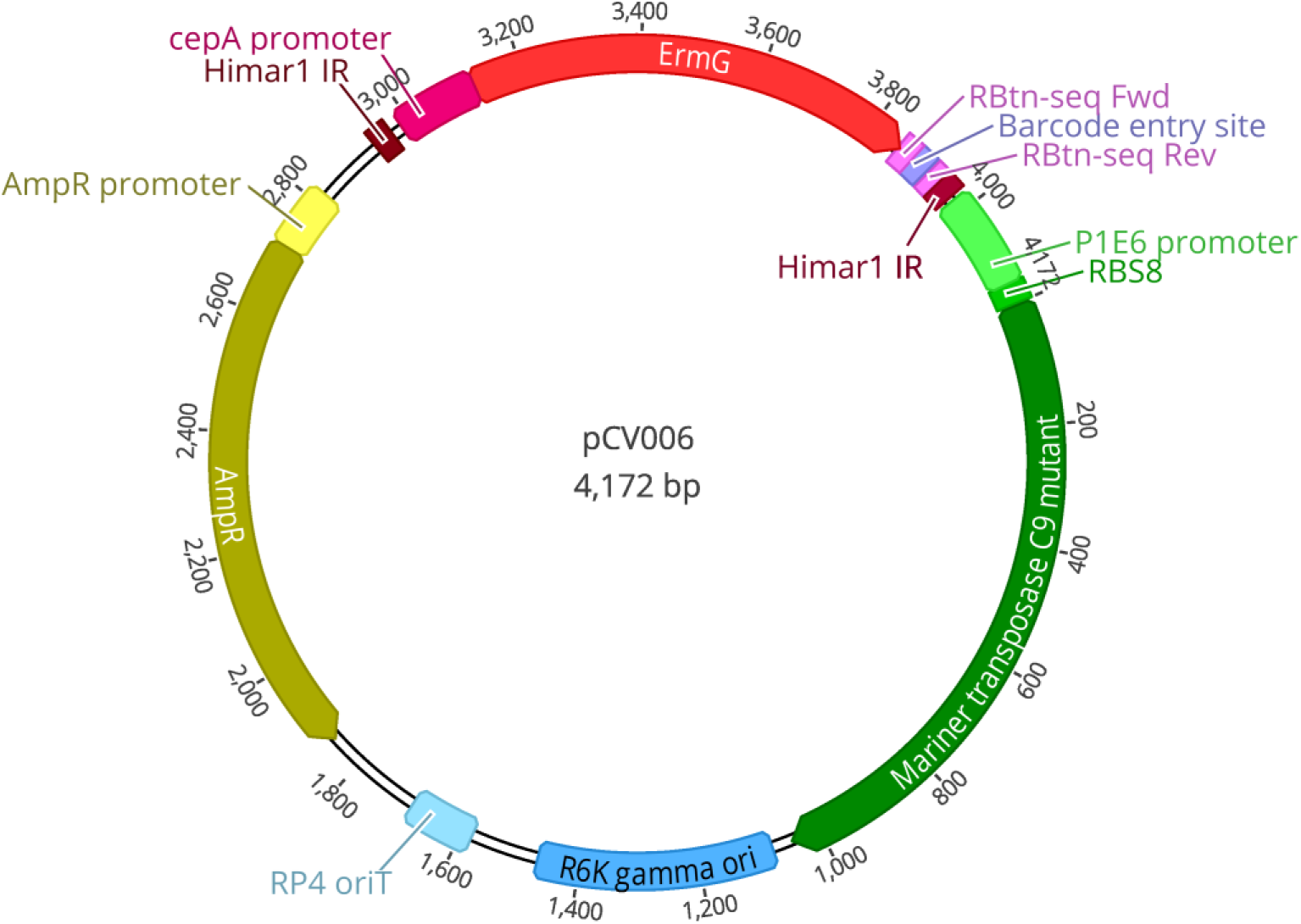

**Supplementary file 1:**
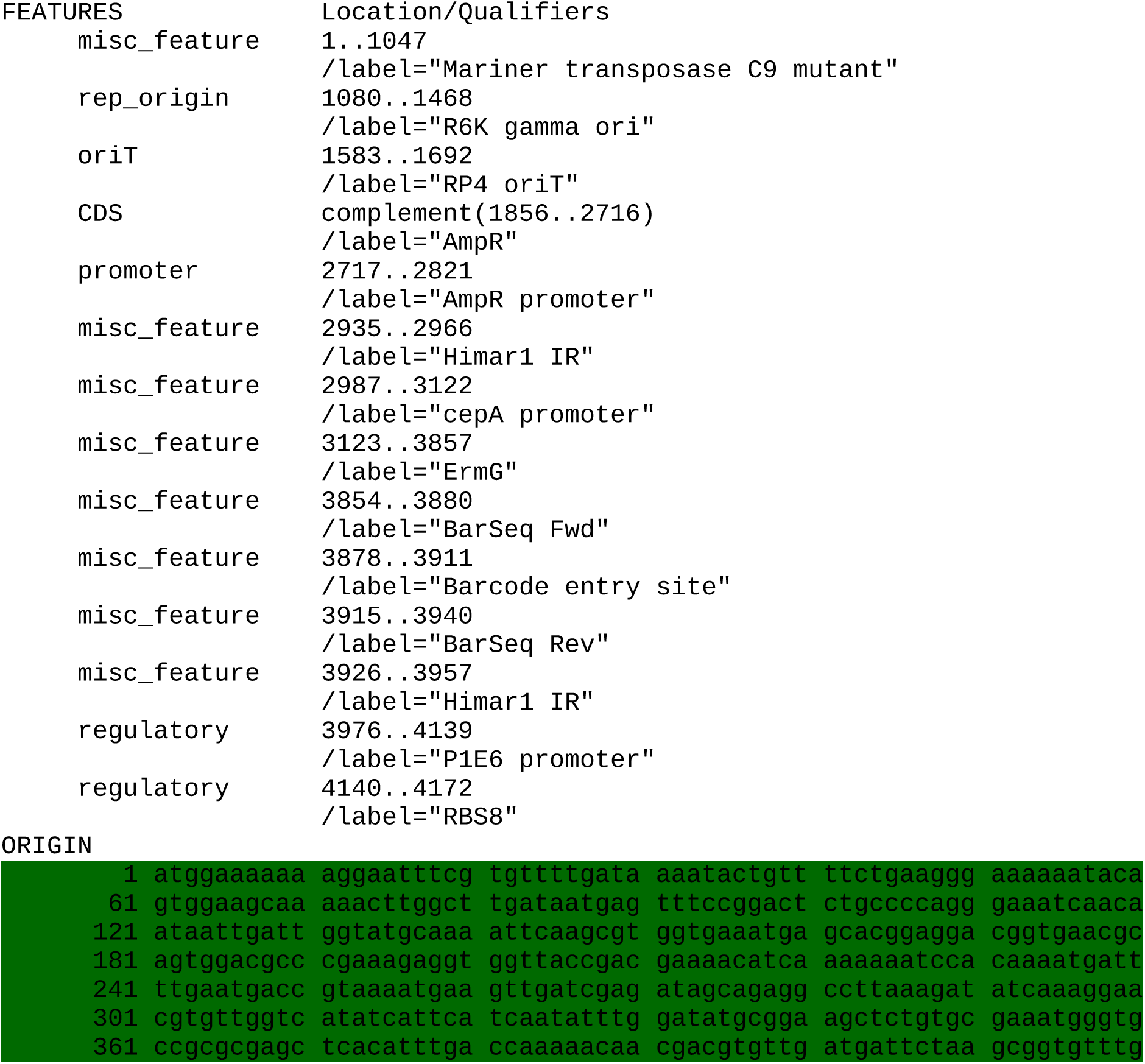
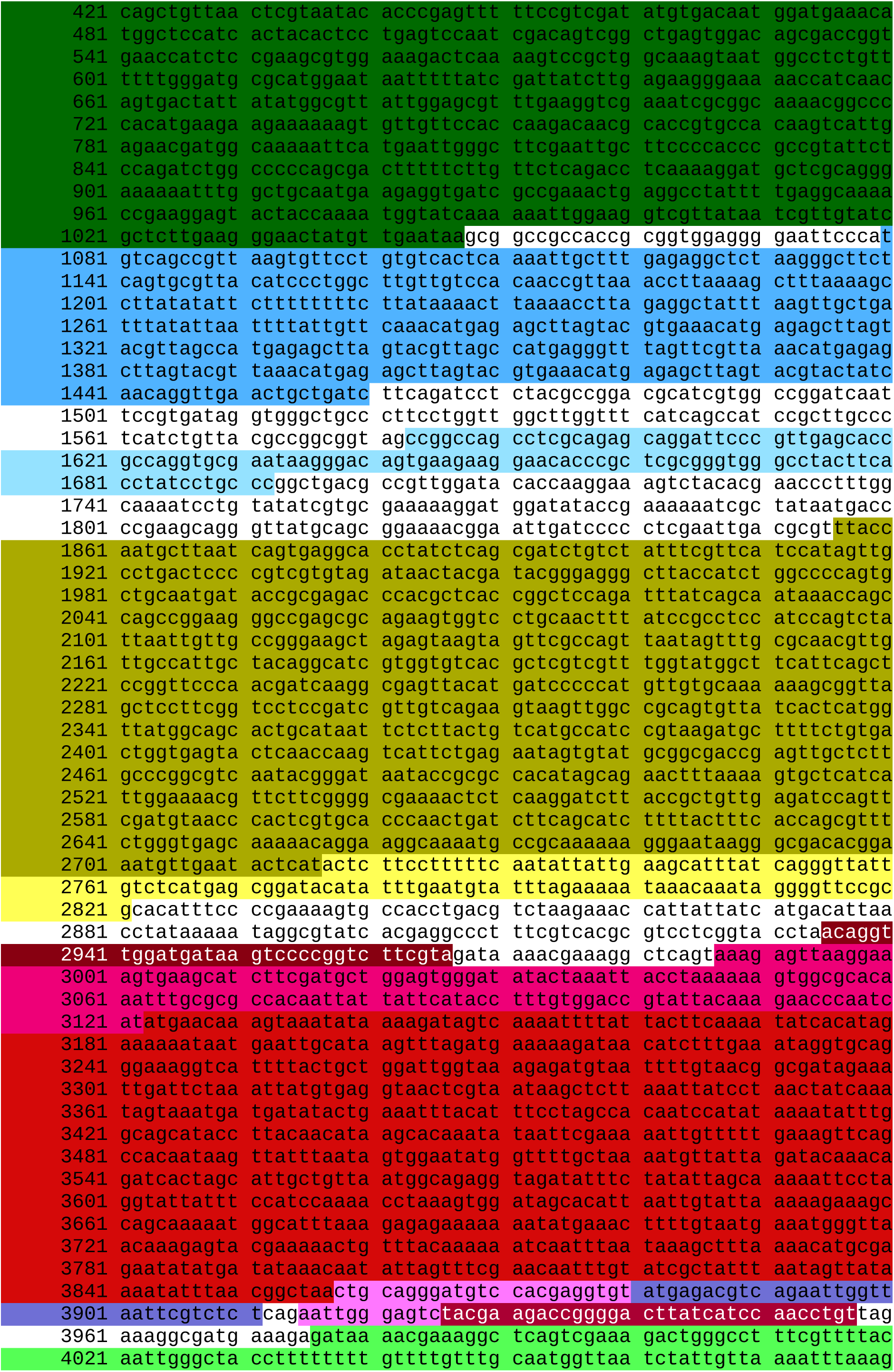
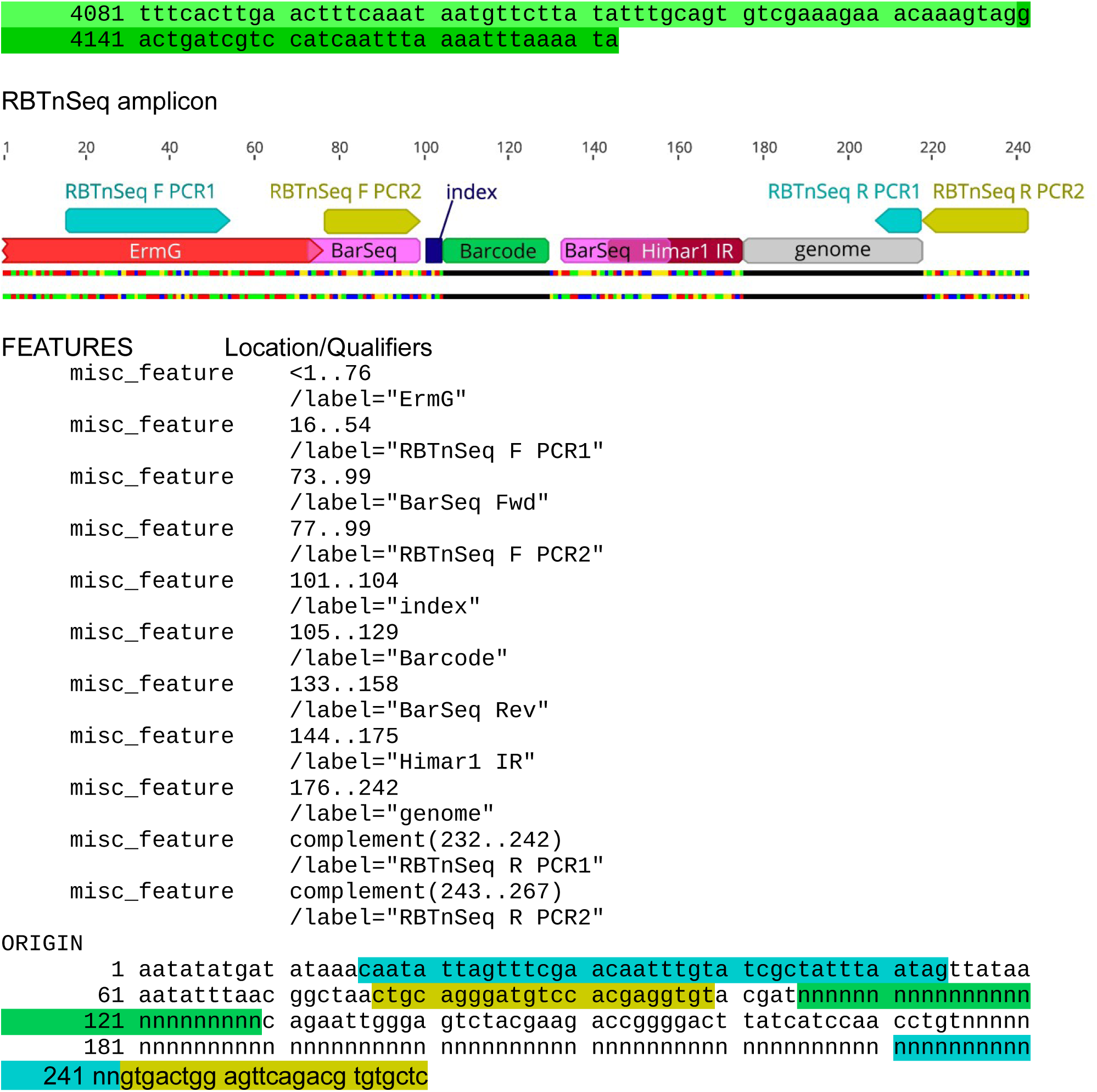
Plasmid features and sequence.

